# Morphometric analysis reveals that the chick cranial neural tube expands as an active shell

**DOI:** 10.64898/2026.05.18.726048

**Authors:** Nimesh Chahare, Chieko Imamura, Nandan L. Nerurkar

## Abstract

Embryonically, the vertebrate brain begins as an approximately uniform, fluid-filled epithelial tube that undergoes rapid volumetric expansion and regionalization to form the morphologically distinct primary brain vesicles. Hydrostatic pressure from fluid secretion into the inner lumen generates tension in the neural tube that has been implicated as a potential driver of cell proliferation during these early stages of brain development. However, a quantitatively rigorous view of 3D morphology and cellular proliferation has remained elusive. Here, we provide a standardized mapping for the mechanical and biological landscape of the developing neuroepithelium along anatomical axes. Using this 3D morphometric framework in chicken embryos, we show that localized curvature characterizes compartmental boundaries. While rapid inflation would typically be expected to stretch and thin the epithelium, we find the opposite: global expansion is coupled with significant tissue thickening, identifying the early brain as an active shell. Moreover, spatial patterns of thickness remain invariant to local curvature. Our results demonstrate a decoupling of geometry and growth, showing that spatially stable distributions of tissue thickness and mitotic activity are maintained throughout massive volumetric expansion, independent of the dramatic geometric reorganization driven by luminal pressure. We conclude that, while tension in the neuroepithelium may contribute to proliferative growth at some level, biological pre-pattern likely plays a driving role in the regionalized expansion of the early embryonic brain.

**Why it matters:** The embryonic brain begins as a simple fluid-filled tube that undergoes rapid and heterogeneous expansion to set up the basic organizational plan of the adult brain. Errors in this process are linked to severe neurological and congenital disorders. This work investigates the biophysical basis of expansion and regionalization of the early brain, a complex three-dimensional process driven by inflation from internal fluid pressure together with active cell behaviors that ultimately produce regionally distinct growth and curvature profiles amid a complex mechanical landscape.

**Graphical Abstract:** 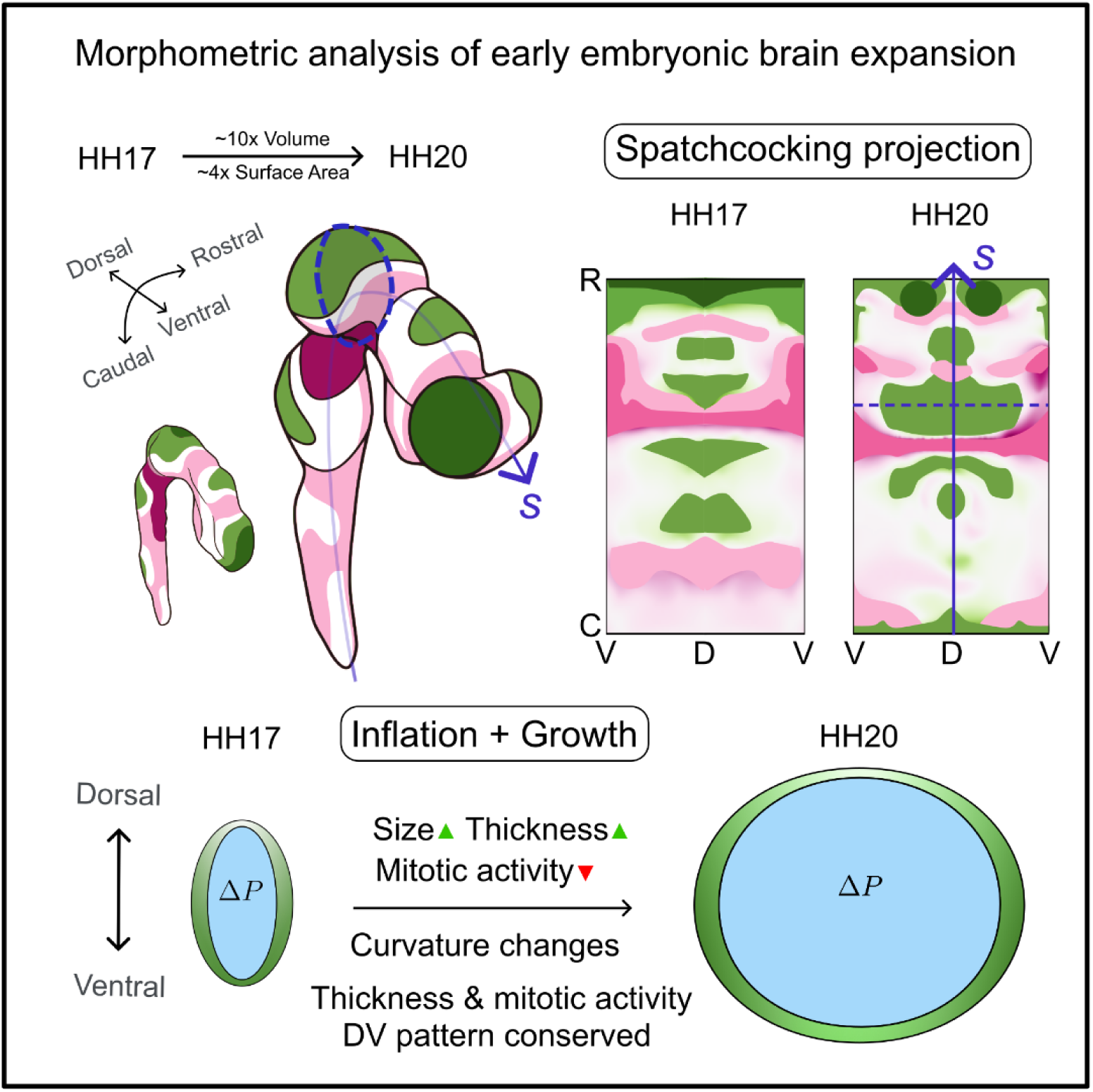

## Introduction

During embryogenesis, the brain originates as a simple, largely uniform epithelial cylinder, the cranial neural tube. The transition of this initial geometry into the primary brain vesicles is a dramatic physical transformation where the neuroepithelium actively deforms and grows to establish the basic organizational plan for the profoundly complex networks that define the adult brain. Errors in early morphogenesis of the cranial neural tube have severe clinical consequences. While gross defects lead to conditions such as anencephaly, more subtle disruptions in early cellular organization are linked to neurodevelopmental disorders (1–3). Central to the early expansion and regionalization of the brain is establishment of a closed, fluid-filled system. In the chick embryo, this occurs via collapse of the neural tube lumen posterior to the hindbrain, separating the enclosed ventricle of the brain forming region from the posterior neural tube, which remains open to the extraembryonic environment until later stages (4). Upon enclosure of the cranial neural tube lumen, cells of the neuroepithelium begin secreting embryonic cerebrospinal fluid (eCSF) into the ventricular space, generating an internal hydrostatic pressure that correlates with a period of rapid volumetric expansion. It has been proposed that intraluminal pressure from eCSF secretion generates tension in the surrounding neuroepithelium that may increase mitotic rates to produce mechanically-driven growth (5–9). While mechanical forces are known in other epithelial systems to drive either passive tissue thinning or active proliferation (10, 11), whether similar mechanisms operate in the embryonic brain has been hindered by a lack of 3D context. Traditional 2D sectioning and isolated explant models inherently strip away the intact geometry of the tube (9, 12), obscuring the spatial relationship between local tissue architecture and the physics of global expansion. Consequently, how 3D morphology and global tissue growth are mechanically coordinated across the emerging domains of the forebrain, midbrain, and hindbrain remains largely unknown. While biochemical signaling is a well-established driver of regionalization (13, 14), the mechanics of this process remain far less understood.

To uncover how this mechanical coordination is achieved, we focused on the chick embryo during a critical 24-hour window of substantial volumetric expansion (HH17–HH20). We developed a 3D morphometric framework that projects the intact 3D neuroepithelium onto 2D anatomical axes, providing a continuous map of local tissue properties. By directly correlating local tissue thickness with local curvature, we find that the embryonic brain diverges from the behavior of a passive elastic shell. Rather than passively stretching and thinning under intraluminal pressure, expansion is coupled with significant tissue thickening. Crucially, we find that the tissue maintains a robust dorso-ventral distribution of both thickness and mitotic activity despite global rounding. Ultimately, our analysis reveals a fundamental uncoupling of tissue architecture from the mechanics of global expansion, ensuring stable compartmentalization during rapid growth.

## Results

### Cranial neural tube enlargement is accompanied by significant lumen rounding

To establish a quantitative baseline for the normal development of the neural tube, we analyzed chicken embryos at stages HH17 and HH20 (Fig. 1a). This 24 hour window represents a period of rapid volumetric expansion (5, 8), yet precedes the peak of neurogenesis when increased cellular complexity arises (15–17). By capturing the system in this state, we can isolate the coordination of global geometry and regional compartmentalization (13) before the emergence of the differentiated mantle layer.

**Figure 1.**
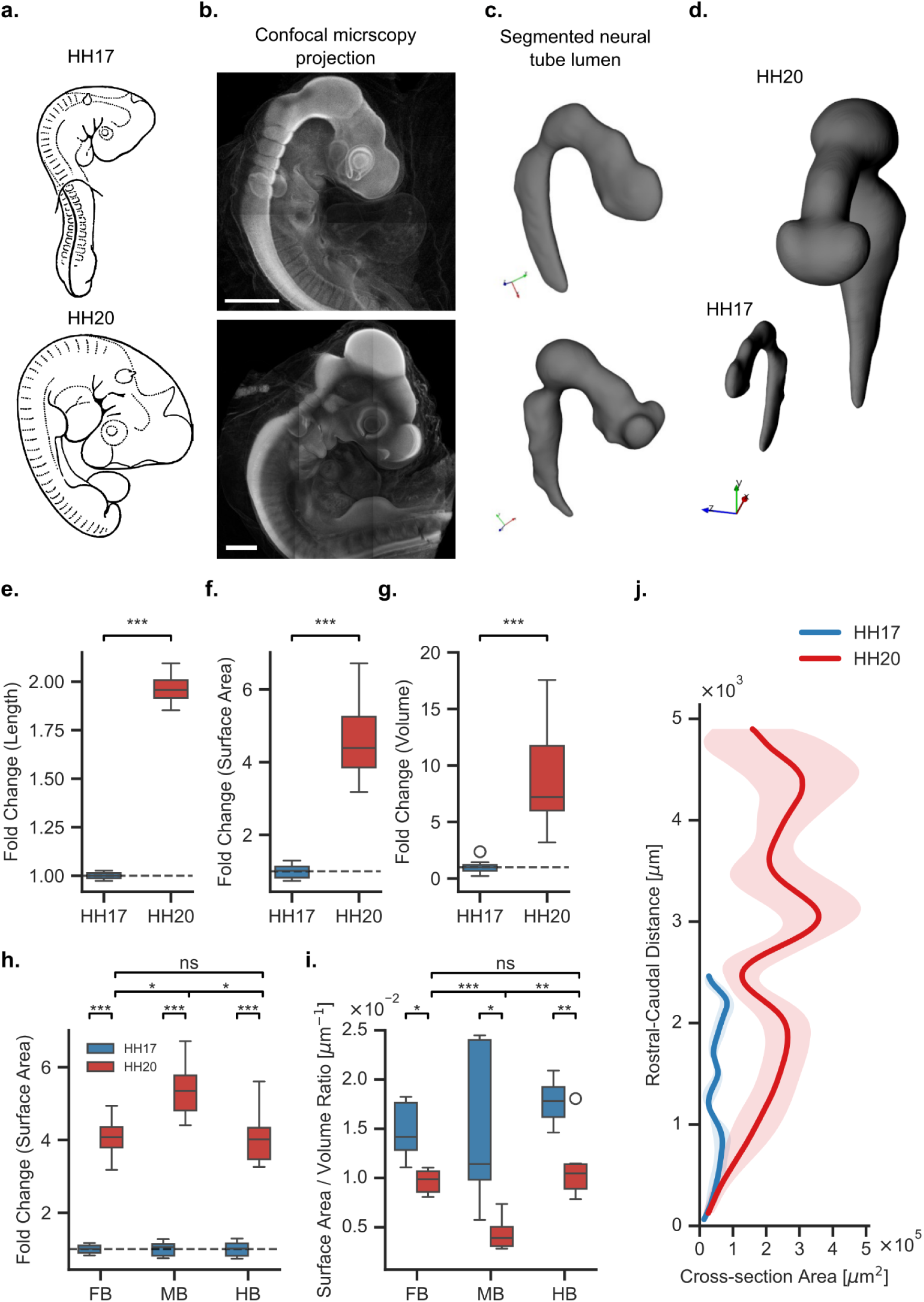
Cranial neural tube enlargement is accompanied by significant lumen rounding. (a) Schematic illustrations of chick embryos at stages HH17 (top) and HH20 (bottom). (b) Maximum intensity projections of optically cleared chick embryos imaged by laser-scanning multiphoton confocal microscopy at HH17 (top) and HH20 (bottom); note dissimilar scales (scale bar = 500 µm) due to size increase between these stages. (c) Segmented 3D meshes of the cranial neural tube lumen at HH17 (top) and HH20 (bottom). (d) Side-by-side comparison of segmented lumen meshes at HH17 and HH20 shown at the same scale to visualize volumetric expansion of the cranial neural tube across this developmental window. (e–g) Fold changes in lumen length (e), surface area (f), and volume (g) between HH17 and HH20. (h) Regional fold changes in surface area for the forebrain (FB), midbrain (MB), and hindbrain (HB). (i) Surface area-to-volume (SA/V) ratio for each compartment at HH17 and HH20. (j) Cross-sectional area profiles along the rostral-caudal axis at HH17 (blue) and HH20 (red). Peaks correspond to primary brain vesicles and valleys to inter-vesicle boundaries. Shaded regions indicate standard deviation across embryos (n = 7 embryos per stage). *p < 0.05, **p < 0.01, ***p < 0.001, ns = not significant.

Because nuclei are densely packed throughout the cranial neural tube, embryos were stained with DAPI as a nuclear stain to broadly visualize tissue boundaries. Samples were optically cleared via a modified 3DISCO protocol (18), and full 3D geometry of the developing brain was imaged using laser-scanning multiphoton confocal microscopy (Fig. 1b). The 3D neuroepithelium and inner lumen were segmented using local thresholding followed by manual refinement to generate triangular meshes. Through this 3D lumen segmentation, we observed a dramatic transition from a relatively uniform tube to a highly compartmentalized and inflated structure (Fig. 1c, d).

Our analysis reveals that this developmental window is defined by substantial physical expansion of the lumen. Between HH17 and HH20, lumen growth is characterized by a doubling of length (Fig. 1e), a four-fold increase in surface area (Fig. 1f), and a nearly ten-fold increase in volume (Fig. 1g). Notably, regional fold-changes indicate that growth is allometric, with the midbrain outpacing forebrain and hindbrain in surface area and volumetric expansion (Fig. 1h, Fig. S1). This geometric transition is most clearly captured by the surface-to-volume (SA/V) ratio, which decreases significantly across all regions from HH17 to HH20 (Fig. 1i). A reduction in SA/V ratio is consistent with transformation from a flattened tube toward a more ballooning geometry, where internal lumen expansion significantly outpaces surface growth to inflate the neuroepithelium. This reduction in SA/V is most pronounced in the midbrain, which reaches the lowest SA/V ratio of the three vesicles. This indicates that the midbrain undergoes the most substantial geometric rounding during expansion—a process likely driven, at least in part, by the accumulation of intraluminal fluid and internal hydrostatic pressure (5, 8).

To move beyond compartmental averages and understand expansion along the rostral-caudal axis, we extracted the medial axis of the curved cranial neural tube using iterative Moving Least Squares (MLS) smoothing (Fig. S2). At both HH17 and HH20, the resulting cross-sectional area profiles reveal a series of peaks and valleys, where peaks correspond to the primary brain vesicles and valleys represent the forebrain-midbrain and midbrain-hindbrain boundaries (Fig. 1j). By HH20, these peaks and valleys become markedly more prominent, consistent with increased compartmentalization and the formation of distinct morphological boundaries as the brain vesicles mature. These data demonstrate that extensive neuroepithelial expansion is coordinated with geometric rounding to establish the compartmentalized structure of the early embryonic brain.

### Standardized 2D mapping enables direct comparison of local neuroepithelium properties across developmental stages

The expanding neural tube presents a non-uniform 3D geometry characterized by the intricate biology of signaling gradients and heterogeneous cell populations (19, 20). To establish a tractable analytical framework for resolving regional mechanical states and local tissue properties, we approach the tissue first as a mechanics and kinematics problem, abstracting the neuroepithelium into a pressurized, thick-walled shell with spatially varying thickness and curvature. This abstraction allows us to resolve the physical state of the tissue, defined by internal tension and hydrostatic pressure, before correlating these mechanical features with biological heterogeneities.

We consider the rostral-caudal (RC) and dorso-ventral (DV) anatomical axes as the primary axes of variation for all localized properties (Fig. 2). To overcome variability in size and shape between individual embryos and across developmental stages, we built upon existing tissue cartography techniques to develop a specialized coordinate mapping framework that standardizes the complex biological manifold of the cranial neural tube (21–23). By mapping the neural tube onto a normalized curvilinear coordinate system and projecting these manifolds into a standardized space, we can decouple intrinsic biological patterns from the global geometry of individual specimens. This enables a direct, node-by-node comparison across different embryos and developmental stages. To implement this, we represented the neural tube as a discrete computational mesh, where every vertex is assigned specific scalar properties (Fig. 2). Probing both tissue morphology and cell behaviors through 3D confocal imaging, this system enables integration of mechanics and biology within a single, unified framework; each vertex can simultaneously represent geometric and mechanical data (e.g., local curvature, tissue thickness, or calculated stress) alongside molecular and cellular data (e.g., gene expression or local mitotic rates). Because these properties vary non-uniformly across the highly curved 3D manifold, a standardized framework is essential for identifying conserved patterns associated with morphogenesis. We achieved this via a 3D-to-2D geometric transformation termed “*spatchcocking*,” implemented using custom Python scripts and the Vedo library (24) to do so (Fig. 2). Borrowing from the culinary technique of splitting and flattening a bird along its midline, this process standardizes the complex 3D architecture into an anatomical coordinate system through a series of discrete mappings. Using the longitudinal medial axis extracted via MLS smoothing, we established a reference frame by defining anatomical dorsal points to align the medial axis relative to the embryo’s dorso-ventral orientation. Local cross-sectional planes, generated perpendicular to this axis, established a localized curvilinear coordinate system used to remap the 3D mesh into a standardized cylindrical frame (Fig. 2). By unwrapping this remapped geometry into a 2D plane and normalizing the rostral-caudal length (from 0 to 1), we generated standardized heatmaps that enable direct, side-by-side spatial comparisons. These heatmaps project biological and mechanical patterns onto a comparable landscape, exposing consistent features otherwise obscured by variations in 3D shape and size. This common coordinate system provides the basis for quantifying spatial variations in local tissue properties across developmental stages below.

**Figure 2.**
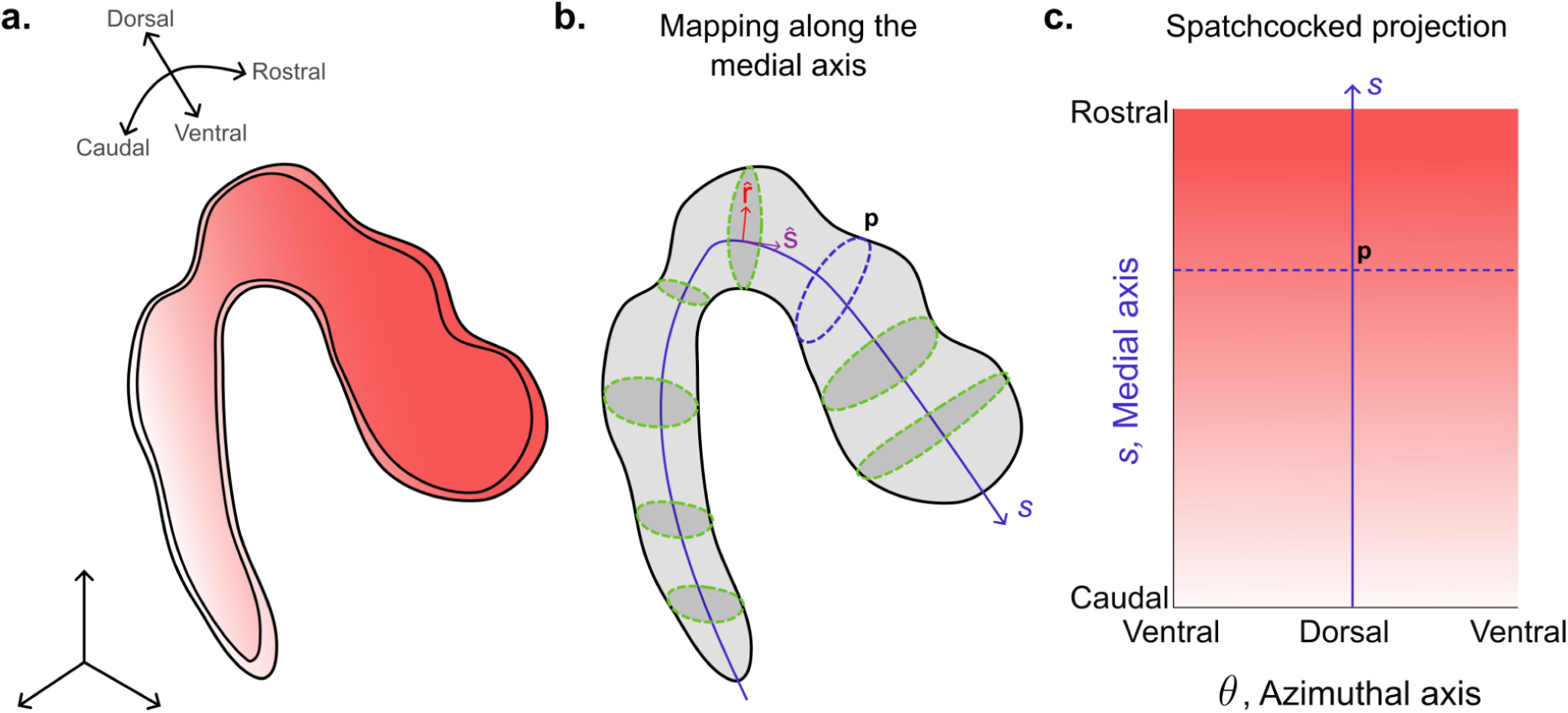
Standardized 2D mapping of the cranial neural tube onto anatomical coordinates. **(a)** Schematic of the cranial neural tube illustrating the anatomical orientation axes: dorsal-ventral and rostral-caudal. The gradient shading represents an arbitrary scalar field mapped continuously across the 3D surface. **(b)** The medial axis (blue curve, *S*) is extracted along the rostral-caudal length of the neural tube. Local cross-sectional planes (green dashed ellipses) are generated perpendicular to this axis, defining a curvilinear coordinate system used to remap the 3D mesh into a standardized cylindrical frame. The azimuthal coordinate *θ* is defined relative to the anatomical dorsal points (*P*) at each cross-section. **(c)** The cylindrical projection is unwrapped into a standardized 2D heatmap, with the medial axis coordinate *s* along the rostral-caudal axis (rostral at top, caudal at bottom) and the azimuthal angle *θ* along the dorso-ventral axis (dorsal at center, ventral at either side). This projection enables direct, node-by-node comparison of local tissue properties across embryos and developmental stages.

### Surface curvature mapping reveals that brain compartmentalization is defined by localized constrictions amidst global inflation

To investigate neuroepithelial rounding as a first application of our standardized coordinate framework, we generated localized maps of surface curvature spanning the entire cranial neural tube from the anterior neuropore to the spinal cord. We implemented a vertex-wise calculation pipeline where local curvature is determined by fitting a quadratic surface to the neighborhood of each vertex on the basal mesh, allowing for the derivation of maximum and minimum principal curvatures (𝑘1 and 𝑘2, respectively), and by extension Gaussian (𝐾 = 𝑘1 𝑘2) and Mean (𝐻 = (𝑘1 + 𝑘2)/2) curvatures, independent of underlying mesh density (Fig. S3).

Gaussian curvature maps reveal a highly organized topological landscape that evolves from HH17 (Fig. 3a, b) to HH20 (Fig. 3d, e) and is remarkably conserved across individual embryos of a given stage (Fig. S4). By pooling data from multiple specimens into binned 2D heatmaps, we identified stereotyped gradients of Gaussian curvature at HH17 (Fig. 3c) that become sharper by HH20 (Fig. 3f). By HH20, prominent dorso-ventral bands of negative 𝐾 were apparent, and align with known anatomical boundaries, including the midbrain-hindbrain and diencephalon-midbrain boundaries (Fig. 3f). These regions of negative Gaussian curvature correspond to anticlastic (saddle-shaped) geometries, indicating where the tissue is locally constricted to maintain compartmental borders. Notably, the fact that the bands of negative 𝐾 become more prominent and sharply defined from by HH20 suggests that compartmentalization is actively reinforced, locally opposing global expansion. In contrast to these boundary constrictions, the centers of the midbrain and forebrain vesicles are characterized by broad patches of positive Gaussian curvature. These synclastic (dome-like) regions where the tissue bulges outward, may represent zones of active inflation. Across all vesicles and stages, dorso-ventral axis profiles of average Gaussian curvature are mediolaterally symmetric relative to the dorsal midline (Fig. 3g). At HH17, the neural tube features sharp curvature gradients, most notably in the midbrain. By HH20, these profiles flatten and smooth as vesicles undergo global rounding. In the forebrain, this shift is marked by localized lateral peaks reflecting emerging cerebral hemispheres and regional bulging.

**Figure 3.**
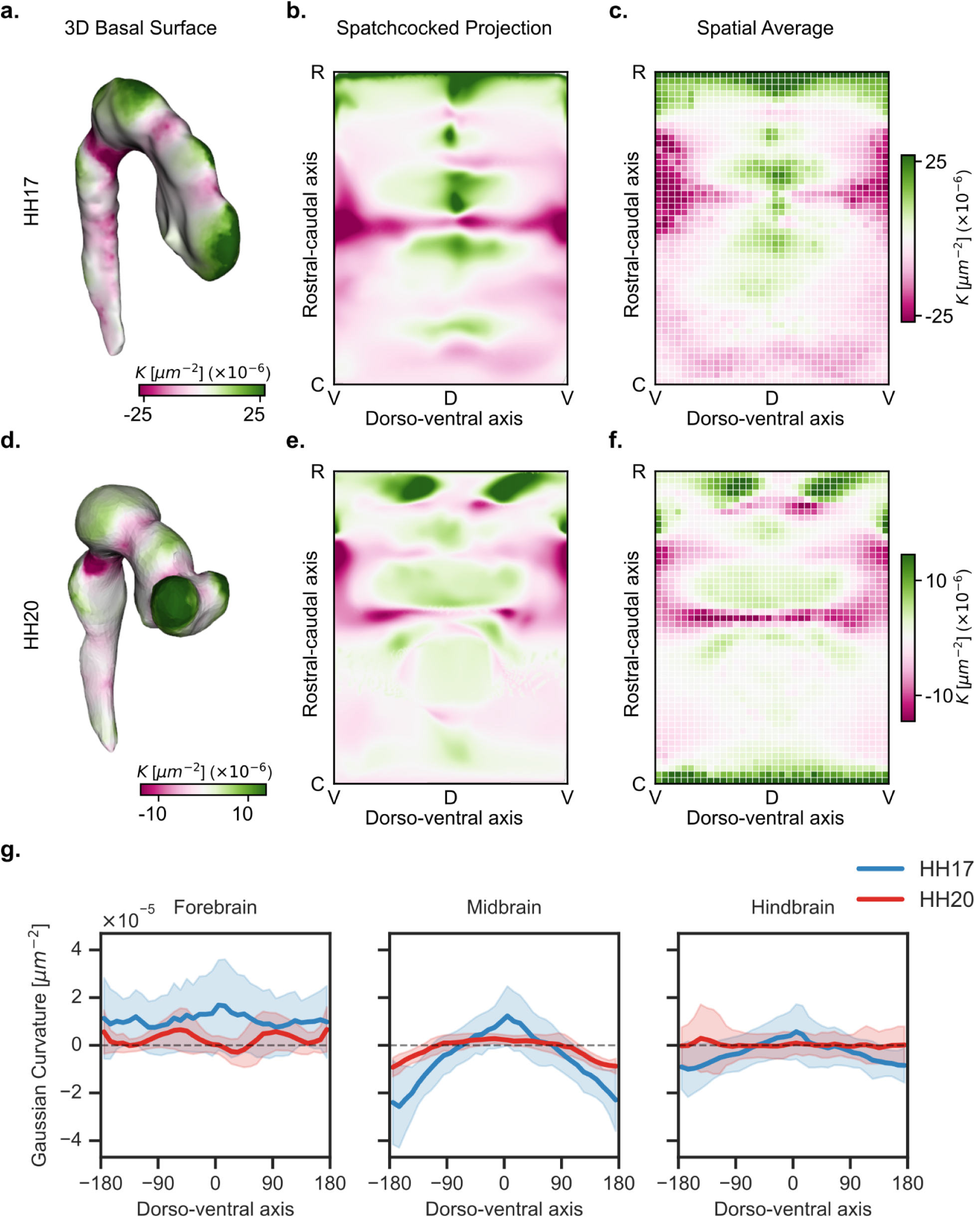
Surface curvature mapping reveals that brain compartmentalization is defined by localized constrictions amidst global inflation. **(a, d)** Gaussian curvature (*K*) mapped onto the 3D basal surface of the cranial neural tube at HH17 (a) and HH20 (d). Green indicates positive (synclastic) curvature; magenta indicates negative (anticlastic) curvature. **(b, e)** Spatchcocked 2D projections of Gaussian curvature for representative embryos at HH17 (b) and HH20 (e), mapped onto the standardized rostral-caudal (R–C) and dorso-ventral (D–V) coordinate system. **(c, f)** Spatially averaged heatmaps of Gaussian curvature pooled across all embryos at HH17 (c) and HH20 (f). **(g)** Dorso-ventral profiles of average Gaussian curvature averaged within the forebrain, midbrain, and hindbrain at HH17 (blue) and HH20 (red). Shaded regions indicate standard deviation across embryos. Dashed line marks 𝐾 = 0. The dorsal midline (0°) is centered, with ventral (±180°) at either side (n = 4 embryos per stage).

To resolve the mechanical state of the expanding neuroepithelium, we mapped the distribution of Mean curvature (𝐻) across the standardized manifolds (Fig. 4a, b, d, e). In pressurized shells, in-plane tension is inversely proportional to mean curvature (σ ∝ 1/𝐻); therefore, the spatial distribution of 𝐻 serves as a proxy for the underlying tension landscape (25, 26). At both HH17 and HH20, 𝐻 remains positive (Fig. 4b, e) and across all regions of the cranial neural tube. A striking transition is visible when comparing the dorso-ventral distribution of Mean curvature between stages. At HH17, the curvature profile is highly non-uniform, characterized by sharp gradients and prominent peaks at the dorsal and ventral midlines (Fig. 4b, g). This identifies the early neural tube cross-section as a pinched ellipsoid where mechanical stress is likely concentrated along the sharpest points of the cross-section.

**Figure 4.**
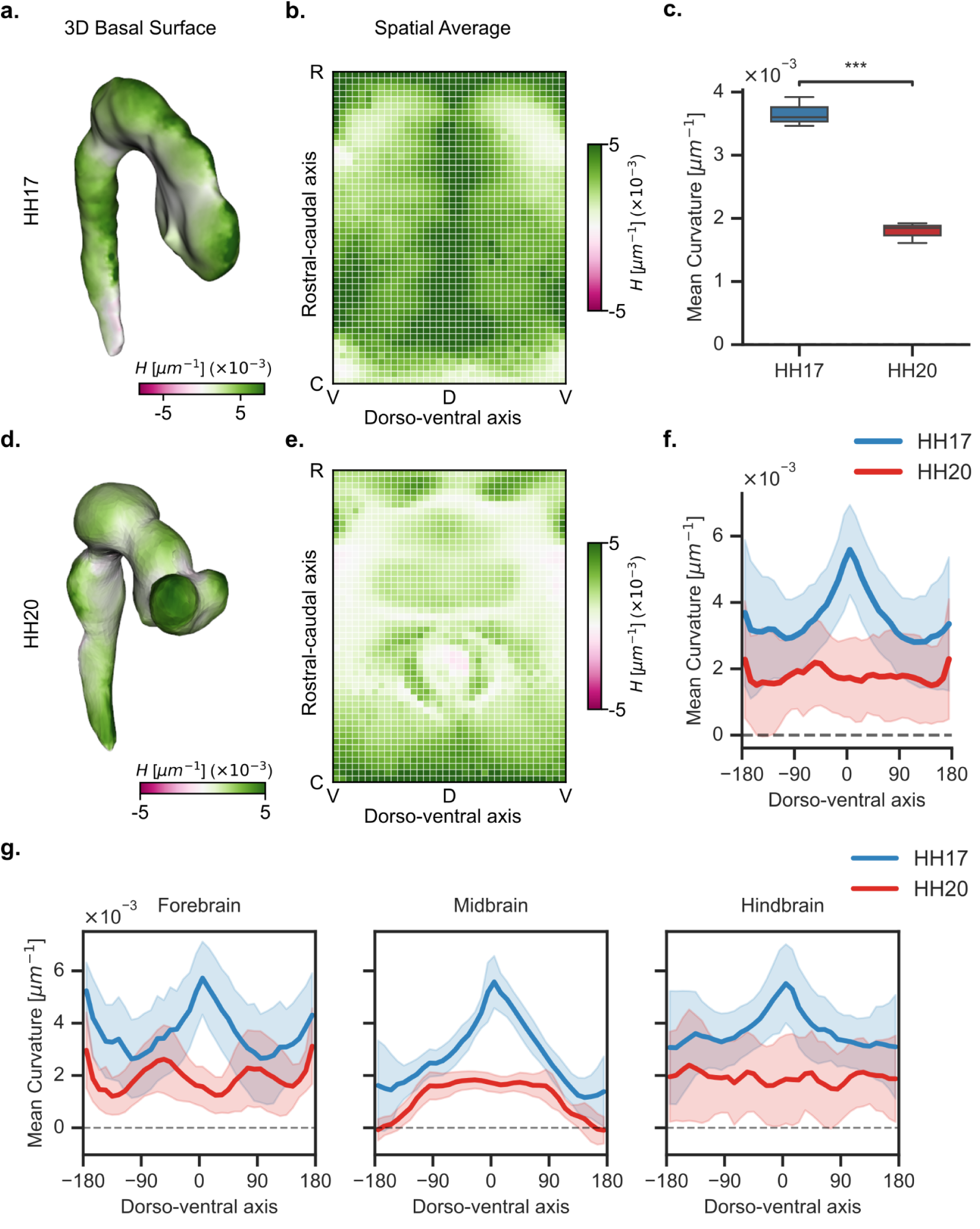
Mean curvature mapping reveals global rounding during volumetric expansion. **(a, d)** Mean curvature (𝐻) mapped onto the 3D basal surface of the cranial neural tube at HH17 (a) and HH20 (d). Green indicates positive curvature; magenta indicates negative curvature. **(b, e)** Spatially averaged heatmaps of mean curvature pooled across all embryos at HH17 (b) and HH20 (e), mapped onto the standardized rostro-caudal and dorso-ventral coordinate system. **(c)** Global mean curvature at HH17 and HH20. **(f)** Dorso-ventral profiles of mean curvature averaged across the entire cranial neural tube at HH17 (blue) and HH20 (red). Shaded regions indicate standard deviation across embryos. **(g)** Regional dorso-ventral profiles of mean curvature for the forebrain, midbrain, and hindbrain at HH17 (blue) and HH20 (red). Shaded regions indicate standard deviation across embryos. n = 4 embryos per stage. *p < 0.001. dorsoventral. Dashed line marks 𝐻 = 0. 𝐻, mean curvature; R–C, rostrocaudal; D–V, dorsoventral. Dashed line marks = 0.

While these patterns define the early geometry, the absolute magnitude of 𝐻 decreases significantly as development progresses (Fig. 4c). As the vesicles undergo a ten-fold increase in volume, the surface becomes locally flat as it becomes globally rounder, a transition clearly captured by the shift in Mean curvature profiles along the dorso-vental axis (Fig. 4f). By HH20, the sharp peaks at the dorsal and ventral midlines effectively flatten, particularly within the midbrain (Fig. 4f). This shift toward a uniform curvature profile indicates that the tissue has rounded into an inflated state, potentially producing a more even distribution of tension across the vesicle circumference. However, it is important to note that a transition toward constant mean curvature does not imply geometric uniformity; rather, it can be achieved through the compensatory variation of principal curvatures as the brain vesicles grow. This is most evident in the forebrain at HH20, where localized peaks in 𝐻 correspond to the emerging cerebral hemispheres (Fig. 4g, lateral peaks), highlighting how regionalized growth persists within the globally expanding shell, consistent with the patterns observed in Gaussian curvature (Fig. 3g). Collectively, these results show that the neural tube does not expand as a simple, uniform cylinder passively stretched by inflation. Instead, the expansion is compartmentalized: while the midbrain and forebrain bulge outward into rounded vesicles, the boundaries between them remain constricted. This reorganization suggests that the neuroepithelium is not a passive shell, but a dynamic manifold where expansion is likely mediated by heterogeneous growth, acto-myosin activity, and material properties.

### The cranial neural tube maintains a dorso-ventral thickness gradient during global enlargement

While surface curvature describes the geometric outcome of inflation, the mechanical response of the neuroepithelium is determined in part by its internal structure. In a pressurized thick-walled shell, local stress is governed by the relationship between curvature (𝐻) and wall thickness (t). We calculated local tissue thickness as the Euclidean distance from the apical to the basal surface, storing the values as vertex-wise attributes on the basal mesh, resolving regional variations in tissue thickness across the entire cranial neural tube (Fig. 5).

**Figure 5.**
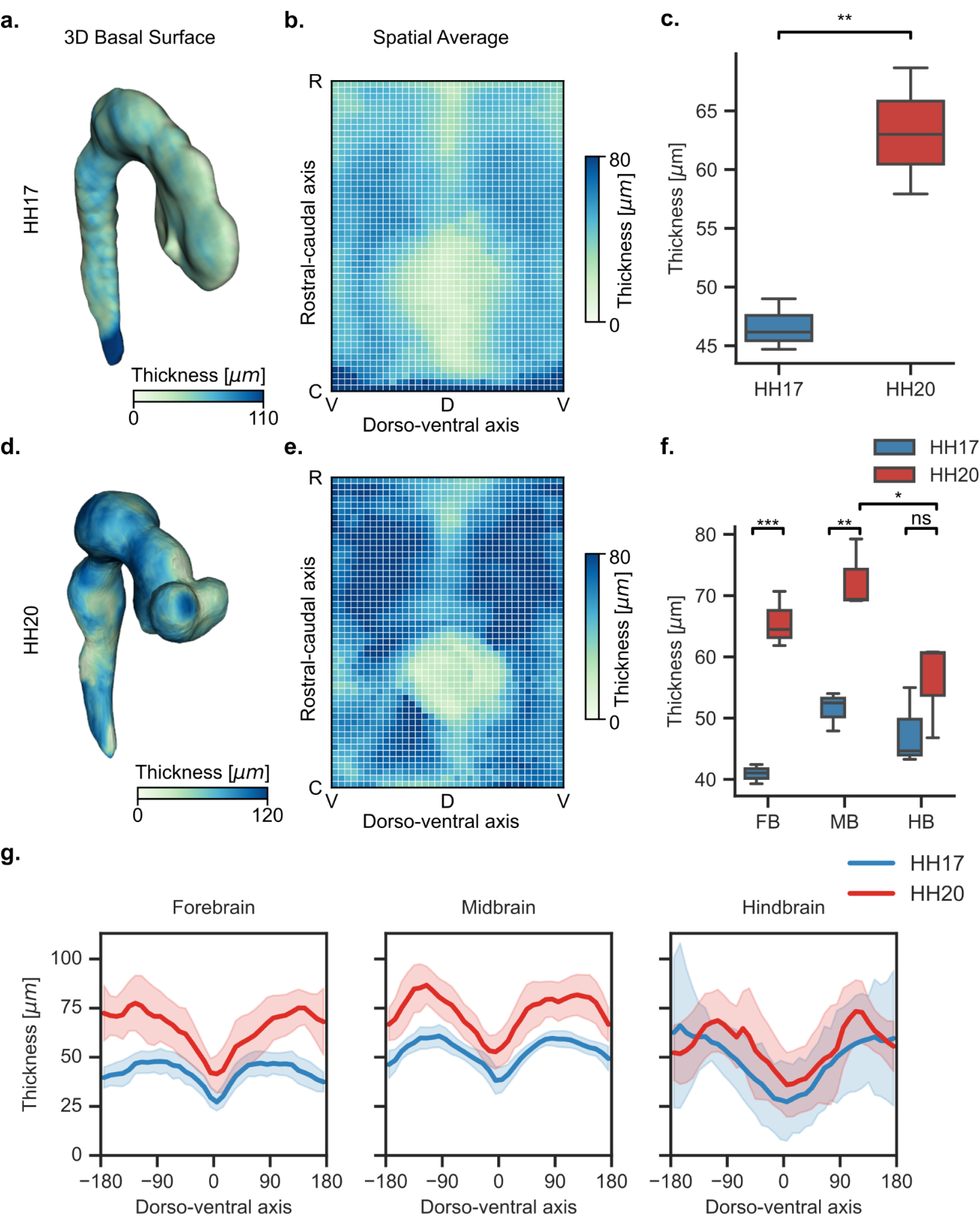
The cranial neural tube maintains a dorso-ventral thickness gradient during global enlargement. **(a, d)** Tissue thickness mapped onto the 3D basal surface of the cranial neural tube at HH17 (a) and HH20 (d). **(b, e)** Spatially averaged heatmaps of tissue thickness pooled across all embryos at HH17 (b) and HH20 (e), mapped onto the standardized rostro-caudal and dorso-ventral coordinate system. **(c)** Global tissue thickness at HH17 and HH20. **(f)** Regional tissue thickness by vesicles at HH17 (blue) and HH20 (red). **(g)** Dorso-ventral thickness profiles for the forebrain (FB), midbrain (MB), and hindbrain (HB) at HH17 (blue) and HH20 (red). Shaded regions indicate standard deviation across embryos. n = 4 embryos per stage. *p < 0.05, **p < 0.01, *p < 0.001, ns = not significant. R–C, rostrocaudal; D–V, dorsoventral.

Our mapping reveals a significant global thickening of the neural tube between HH17 and HH20, with average thickness across the cranial neural tube increasing from 46.5 ± 1.8 µm to 64.0 ± 4.7 µm (Fig. 5c, Fig. S7). This thickening is also apparent via regional variations on the heatmaps (Fig. 5b, e). While the hindbrain roof plate remains the thinnest point on the brain across both stages, the lateral walls of each vesicle thicken markedly (Fig. 5b, e). While all compartments thicken between stages, this growth is non-uniform; the hindbrain exhibits the least increase, remaining significantly thinner than the midbrain by HH20 (Fig. 5f).

Thickness profiles along the dorso-ventral axis confirm that the tissue maintains a heterogeneous distribution across both stages (Fig. 5g, Fig. S6). However, the relative dorso-ventral pattern of apico-basal thickness is conserved across all three primary vesicles, with the forebrain, midbrain, and hindbrain each exhibiting a characteristic profile defined by thick lateral walls and a thin dorsal midline. Notably, while the ventral midline also represents a local thickness minimum in all three compartments, it remains consistently thicker than the dorsal roof plate. The qualitative similarity of these profiles across the rostral-caudal axis suggests that the mechanisms governing thickness distribution may not be vesicle-specific, but rather represent a fundamental organizational principle of the early cranial neural tube.

Strikingly, at HH20, this characteristic dorso-ventral thickness profile is preserved even as the brain rounds out and its mean curvature approaches uniformity (Fig. S5, Fig. S6). Accordingly the strong negative correlation between tissue thickness and curvatures that is apparent at HH17 is lost by HH20 (Fig. S5). In a passive elastic shell with uniform mechanical properties, wall thinning is a geometric consequence of stretching to accommodate surface expansion. However, we find the opposite: thickness increases globally with the characteristic dorso-ventral gradient shifting toward higher absolute values (Fig. 5b, e). To investigate the cellular basis of this thickening, we next quantified the spatial distribution of cell proliferation. By integrating mitotic rates into our 3D framework, we examined whether localized cell division spatially corresponds to the maintenance of these thickness gradients during expansion.

### Spatial patterns of mitotic activity are preserved across global expansion

To map the spatial distribution of cell division, we performed whole-mount immunostaining for phospho-histone H3 (pHH3), a marker of mitotic cells. All pHH3+ cells within the neural tube were identified, and their 3D coordinates used to compute a local density field via fixed-radius neighborhood search (R = 100 µm), which was then transferred onto the mesh using inverse distance weighting interpolation (Fig. 6a, d, Fig. S8a-c).

**Figure 6.**
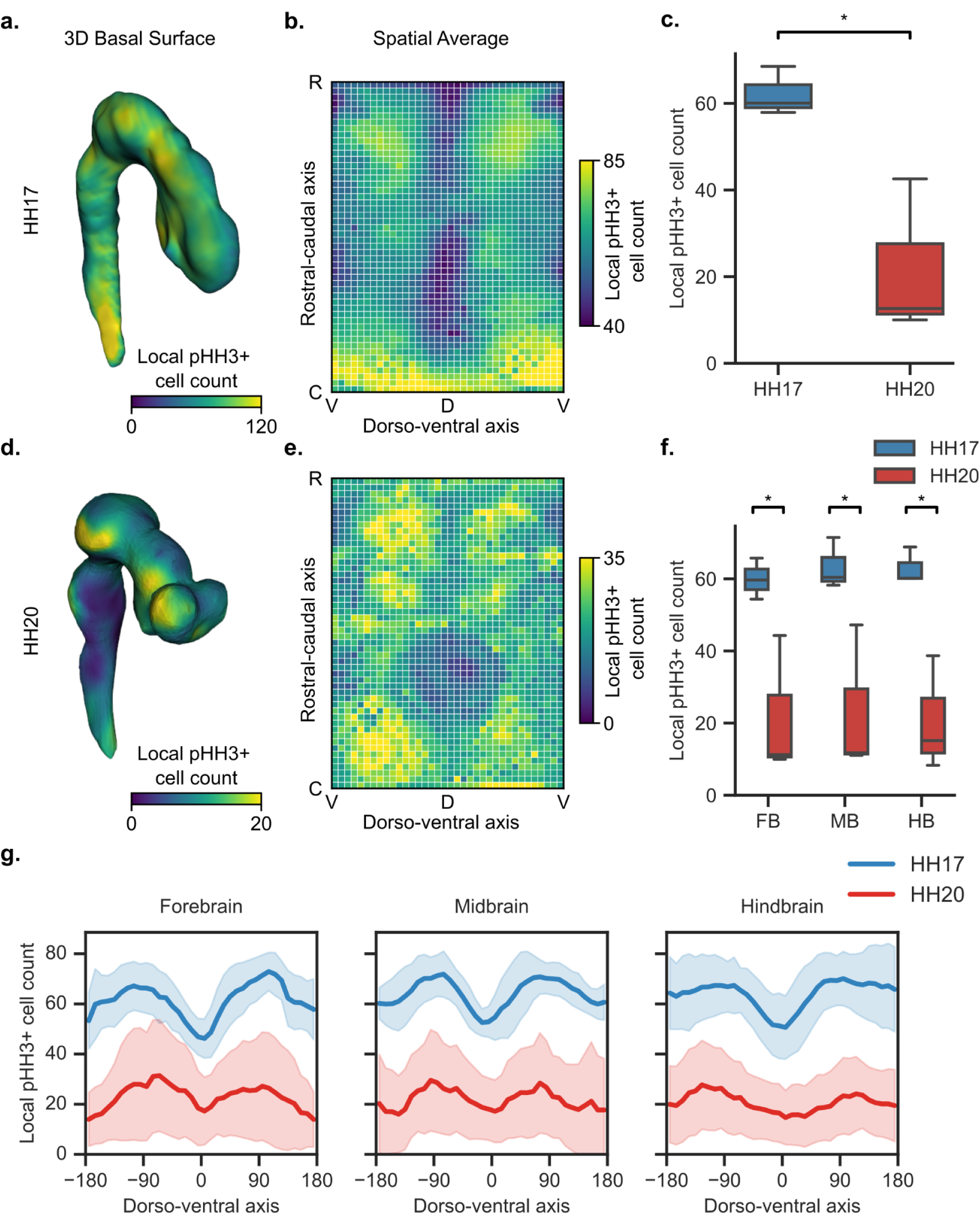
Spatial patterns of mitotic activity are conserved across global expansion. **(a, d)** Local pHH3+ cell density mapped onto the 3D basal surface of the cranial neural tube at HH17 (a) and HH20 (d). Note the difference in color scale between stages. **(b, e)** Spatially averaged heatmaps of local pHH3+ cell counts pooled across all embryos at HH17 (b) and HH20 (e), mapped onto the standardized rostro-caudal and dorso-ventral coordinate system. **(c)** Global local pHH3+ cell counts at HH17 and HH20. **(f)** Regional local pHH3+ cell counts by compartment at HH17 (blue) and HH20 (red). **(g)** Dorso-ventral profiles of local pHH3+ cell counts for the forebrain (FB), midbrain (MB), and hindbrain (HB) at HH17 (blue) and HH20 (red). Shaded regions indicate standard deviation across embryos. n = 3 embryos per stage. *p < 0.05. R–C, rostrocaudal; D–V, dorsoventral.

At HH17, pHH3+ cell counts are elevated along the lateral walls and attenuated at the dorsal and ventral midlines (Fig. 6b, e). Between HH17 and HH20, local pHH3+ counts decrease significantly across all compartments (Fig. 6c, f, Fig. S8g), consistent with the transition from a rapidly proliferative state toward the onset of neurogenic differentiation (15, 27). Despite this global reduction, mitotic activity remains laterally enriched across all vesicles (Fig. 6g).

Ultimately, this dorso-ventral distribution of mitotic activity is conserved across both stages (Fig. 6g, Fig. S6d). Taken together, these results reveal that the early embryonic brain maintains spatially stable distributions of tissue thickness and mitotic activity throughout a period of dramatic geometric transformation. Rather than undergoing passive mechanical deformation in response to luminal pressure, the early neuroepithelium exhibits proportional growth that preserves its established morphological template.

## Discussion

By projecting the complex 3D chick cranial neural tube into a standardized 2D framework, we provide a quantitative map of early brain expansion during key stages of regionalization. These analyses highlight how tightly stereotyped the global morphology of the cranial neural tube is while undergoing dramatic physical transformation, including the precise location and curvatures of vesicle boundaries that are thought to arise through localized patterns of acto-myosin activity (28). How long range coordination this spatial pattern is established is unclear, but unlikely to be fully mediated directly through morphogenic cues, given length scale over which these patterns must be established (∼5 mm). However, the presence of a continuous, fluid-filled lumen may create a mechanical source of long-range communication, as local constrictions would alter global tension through changes in fluid pressure. How such interactions are integrated with localized signaling interactions to establish and maintain the global organization of the cranial neural tube will be important to consider in future work.

As the brain undergoes substantial volumetric expansion (Fig. 1g), we observed that it transitions from an elliptical tube into a highly compartmentalized structure defined by rounded vesicles and localized boundary constrictions (Fig. 3). This morphological shift is most clearly captured by Mean curvature (H) measurements, which reveal a significant flattening of the tissue surface between HH17 and HH20 (Fig. 4c, f). This transition from a constricted tube to a series of rounded vesicles suggests that internal hydrostatic pressure from eCSF secretion may act as a driver of this physical transformation into the primary brain vesicles (8, 29, 30).

Despite this global rounding, however, Gaussian curvature (𝐾) maps reveal a highly organized topological landscape in which dorso-ventral bands of negative 𝐾 at the midbrain-hindbrain boundary (MHB) and diencephalon-midbrain boundary (DMB) persist and sharpen between stages (Fig. 3b, e), indicating compartmentalization is actively reinforced during inflation. These anticlastic zones are consistent with the view that compartmentalization is initiated by differential contractility mediated by circumferential actomyosin fibers at the sulci and subsequently maintained by growth (28, 29).

From the classical engineering perspective, a pressurized vessel thins as it expands, a phenomenon observed in the blastocyst and other lumenized tissues (31–33). Although the present study does not directly measure tissue forces, the neuroepithelium’s thick-walled geometry and spatially varying thickness render simple thin-shell approximations such as the Law of Laplace 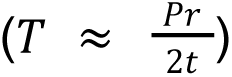 insufficient. A thick-walled shell with spatially varying thickness develops stress concentrations at its thinnest regions alongside internal bending moments. To put this in context, we performed plane-stress finite element simulations of the midbrain cross-section using measured representative geometres of the forebrain, midbrain, and hindbrain (Fig. S9). This proof-of concept model illustrates that stress is concentrated at the thin dorsal regions and varies through the wall thickness, generating internal bending moments that a simple thin-shell approximation cannot capture (Fig. S9). The mechanical expectation is therefore that thinner regions would stretch disproportionately, becoming thinner still. In stark contrast, the neuroepithelium thickens globally while maintaining its dorso-ventral thickness gradient (Fig. 5c). While previous studies have decoupled the elastic and growth components of circumferential expansion (8, 29), radial growth (thickness) has remained largely uncharacterized. Recent evidence suggests thickness is regulated by luminal pressure; pharmacologically increasing pressure maintains thickness during expansion, while pressure reduction induces thickening during lumen collapse (30). We identify radial thickening as a regulated component of expansion that offsets the thinning expected from passive stretching. This active growth maintains the spatial distribution of tissue thickness even as the surface curvature is reorganized by luminal pressure.

A central finding of this study is that the dorso-ventral thickness gradient and laterally peaked mitotic distribution remain spatially invariant despite substantial flattening of the mean curvature profile between HH17 and HH20 (Fig. 4g, 5g, 6g). By HH20, the neuroepithelium contains a mixed population of actively dividing progenitors and cells differentiating into distinct cell fates (15), and so the global reduction in mitotic activity is to be expected; the dorso-ventral mitotic distribution, however, is preserved from HH17 to HH20. Each cell population likely carries distinct mechanical properties, differences in cortical tension, and adhesion that may themselves contribute to the spatially heterogeneous growth response observed here. Indeed, regional variation of material properties has been implicated in differential expansion of the hindbrain roofplate (30). Based on well-known dorso-ventral and rostro-caudal asymmetries in the expression of key morphogens including Hedgehog, BMP, FGF, and Wnt ligands, moving forward it will be important to identify how such orthogonal gradients of morphogenic cues are integrated in 3D to specify material properties, contractility, and growth heterogeneities to establish precisely stereotyped expansion and regionalization.

The dorso-ventral thickness gradient described here, with thick lateral walls and a thin dorsal midline, is consistent with the hindbrain measurements of McLaren et al. (30), and with the dorso-ventral gradient of mitotic activity in the forebrain reported by Garcia et al. (8). The 3D framework introduced here resolves both patterns continuously across all three vesicles simultaneously. As multi-scale high resolution imaging continues to advance, it will be important to link these tissue-scale analyses to individual cell shapes, division orientations, and local cell density variations, all of which are known to be mechanosensitive (10, 34, 35). The spatchcocking framework we present here is designed to be extensible. Integration with techniques such as fluorescent in situ hybridization would allow gene expression patterns to be mapped directly onto the same coordinate system, providing a basis for resolving how molecular gradients and mechanical signals interact across developmental stages and for testing competing anatomical models of early brain organization (19, 36).

Tension on the neuroepithelium, generated by hydrostatic pressure from the secretion of eCSF into the lumen, has been suggested in a handful of studies to regulate neuroepithelial mitotic rates through mechanotransduction pathways including Focal Adhesion Kinase (FAK) activity (4, 6, 8, 37, 38). Although our descriptive analysis here does not directly test a causal link between tension and cell proliferation, there are some important correlative insights to note. The current literature describes a chemomechanical model in which hydrostatic pressure creates a permissive tension state for proliferation, while molecular signals, including FGF8 from the anterior neural ridge and BMP4 from the roof plate, act as regional modulators of this mechanical stimulus (8). Our observation that mitotic activity is laterally enriched and attenuated at the dorsal midline is consistent with this framework: BMP4, secreted at the roof plate, is known to inhibit proliferation at the dorsal midline, while FGF8 and Shh promote growth in complementary domains (19, 39–41). The interaction between biochemical signaling and geometric constraints produces a non-uniform dorsoventral distribution of mitotic activity. Importantly, the spatial stability of this mitotic distribution across a period of substantial geometric reorganization and expected shifts in the mechanical landscape of the neural tube (Fig. S9) suggests that the molecular pre-pattern, rather than the local tension state alone, may be the dominant regulator of how and where cell proliferation is enriched in the cranial neural tube. In a simplified, linear 2-D finite element model with homogeneous material properties, we indeed find that tension is highest in thinner regions of the neuroepithelium, contrasting with our experimental findings that proliferation rates are highest in thicker regions. However, in previous studies, experimental perturbations that reduce and increase intraluminal pressure have shown corresponding changes in cell proliferation, suggesting that under sufficiently large exogenous perturbations, tension does inform mitotic rates in the neuroepithelium (8, 29, 30, 37, 38). Reconciling the role of endogenous tension with cell proliferation may require an integrated model that accounts for 3-D variations in morphogenic cues, passive mechanical properties, and active cell-generated forces, building on the dataset and spatchcocking framework introduced here, as well as key prior studies capturing growth and contractility in a continuum modeling framework (8, 29).

The standardized morphometric framework introduced here provides a continuous, quantitative view of the intact embryonic brain that traditional sectioning approaches cannot capture. Applying this framework reveals that the early neuroepithelium maintains spatially stable distributions of tissue thickness and mitotic activity throughout massive volumetric expansion, independent of the dramatic geometric reorganization occurring over the same period. These results point to a growth program that is preserved robustly through the earliest and most dramatic phase of brain morphogenesis.

## Methods

### Chick embryo collection

Fertilized White Leghorn chicken eggs (University of Connecticut Poultry Farm) were incubated at 37°C and approximately 60% humidity until the stages HH17 (66 hours) and HH20 (90 hours) according to the Hamburger and Hamilton staging system (Hamburger and Hamilton 1951).

### Immunostaining

Embryos were fixed overnight in 4% PFA, washed in PBS, and processed for either whole-mount immunostaining or cryosectioning. For whole-mount staining, embryos were permeabilized in PBTS 1% (1% Triton X-100, 1% DMSO in PBS), blocked in 10% heat-inactivated goat serum, and incubated with primary antibodies overnight at 4°C followed by secondary antibodies overnight at 4°C. For cryosectioning, fixed embryos were moved to 30% sucrose, embedded in OCT, and sectioned at 16 µm on a Leica cryostat. Sections were blocked and incubated with primary and secondary antibodies before mounting with Fluoromount-G. Primary antibodies used in both protocols included rabbit anti-Phospho-Histone H3 Ser10 (1:500; Millipore Sigma 06-570), detected with a Cy3-conjugated goat anti-rabbit secondary antibody (Jackson ImmunoResearch); DAPI (1:1000; Invitrogen) was used as a nuclear counterstain in whole-mount preparations. Following whole-mount staining, embryos were optically cleared using a 3DISCO protocol (18).

### Confocal microscopy

For 3D imaging of the neural tube, samples were imaged using a Zeiss LSM880 laser-scanning confocal microscope equipped with a 10x air objective (NA 0.45). Z-stacks were acquired with a 4.5 µm step size at a resolution of 256 x 256 pixels with each pixel length of ∼5.5 µm, capturing a total of 400–500 slices in the Z-dimension. DAPI was excited at 740 nm using a two-photon laser and captured at 400 nm, while Phospho-Histone H3 (Cy3) was excited at 561 nm and detected at 594 nm via a GaAsP-T1 detector. Individual tiles were stitched using Zeiss ZEN software to generate a composite image of the entire neural tube.

### Neural tube segmentation

All codes to follow the analysis pipeline are available as supplementary material. To segment the neuroepithelium and inner lumen, czi files were first converted to.tiff format and downsampled four-fold using the Scale function in Fiji/ImageJ (42). The DAPI channel was isolated and subjected to a gamma correction of 0.5 to enhance structural contrast. Volumetric segmentation was performed in 3D Slicer (43) using a hybrid approach: an initial mask of the neuroepithelium was generated via local thresholding, followed by manual refinement to ensure anatomical accuracy. The resulting masks were post-processed to fill internal holes and close gaps; all 3D visualization of the processed.tiff files and masks was performed using napari (44). To extract the inner lumen surface, a custom Python script utilizing the tifffile and scipy libraries was employed to isolate the tissue-lumen interface from the processed 3D masks. Finally, these 3D masks were converted into triangular meshes using a marching cubes algorithm and the Trimesh Python library. The reconstructed meshes were exported in.stl and.ply formats to facilitate 3D surface rendering and volumetric analysis.

### Tissue Shape and Morphometric Analysis

For 3D morphological characterization, mesh processing and morphometric analysis were performed using the Vedo Python library (24). To standardize the mesh topology while preserving the anatomical integrity of the neural tube, meshes were first decimated to 200 vertices and subsequently subdivided three times. Surface noise was reduced using a Windowed Sinc smoothing filter, which minimizes the volume shrinkage and feature blurring typically associated with standard Laplacian smoothing.

Local surface curvature of the smoothened mesh was then calculated using a custom Python script. For each vertex on the mesh, a quadratic surface was fitted to a local neighborhood of adjacent vertices, from which the Gaussian and Mean curvatures were calculated. This fitting procedure was iterated across all vertices of the mesh, ensuring a robust and accurate representation of the tissue topography independent of the underlying element size.

Tissue thickness was quantified by calculating the Euclidean distance between the segmented inner (lumen) and outer (basal) surfaces of the neural tube. Using the distance_to function from the Vedo library, a point-to-mesh distance map was generated by computing the distance from each vertex on the lumen surface to the nearest polygonal face of the outer surface mesh. A key advantage of using the Vedo framework is that these calculations result in vertex-wise scalar fields for both curvature and thickness, which are stored directly within the same mesh object to facilitate integrated spatial analysis.

### Proliferation analysis

To quantify the spatial distribution of cell division, Phospho-Histone H3 (pHH3) positive nuclei were identified from confocal image stacks using the Spots detection module in Imaris (Bitplane), with a specified mean spot diameter of 20 µm. The 3D spatial coordinates of detected spots were exported and processed using a custom Python script. To ensure anatomical accuracy, these points were filtered against the 3D neuroepithelium mask, and any spots located outside the tissue boundaries were excluded to form a discrete point cloud. To transform these discrete points into a continuous spatial signal, a local density field was computed using the density() function from the Vedo Python library, which implements a fixed-radius neighborhood search. The local pHH3+ cell count D at any spatial coordinate x was defined as:

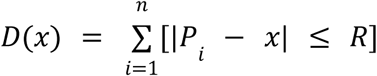

where Pᵢ is the position of the i-th pHH3+ cell and R = 100 µm. This radius was selected to exceed the maximum measured tissue thickness at both developmental stages ensuring the sampling sphere spans the complete apical-to-basal extent of the neuroepithelium at every surface location. The density field was computed over a high-resolution voxel grid matched to the bounding box of each specimen and projected onto the standardized basal mesh using Inverse Distance Weighting interpolation (Shepard’s method, power parameter p = 2), where each mesh vertex was assigned a weighted average of the nearest voxel values. The resulting vertex-wise scalar data were integrated into the standardized coordinate framework to enable direct spatial comparison of mitotic activity across embryos and developmental stages.

### Spatchcocking transformation

To facilitate spatial analysis across varying embryo geometries and developmental stages, a 3D-to-2D geometric transformation, termed “spatchcocking”, was implemented using custom Python scripts and the Vedo library (24). This process standardized the complex 3D architecture into a canonical 2D coordinate system through a series of discrete mappings. Initially, the longitudinal medial axis (Rostral-Caudal axis) of the neural tube was extracted from the 3D lumen mesh using iterative Moving Least Squares (MLS) smoothing. To establish a biologically relevant reference frame, anatomical dorsal points were manually selected on the mesh to define a secondary vector, allowing for the alignment of the medial axis relative to the dorsal-ventral axis of the embryo.

A series of local cross-sectional planes were then generated perpendicular to the computed medial axis. The points of intersection between these planes and the embryo were used to define a localized curvilinear coordinate system, establishing a new set of target coordinates along the Rostral-Caudal and Dorsal-Ventral axes. This transformation was executed via a Thin Plate Spline warping method using the warp() function in Vedo. By providing the original mesh points and their corresponding transformed coordinates as inputs, the function performed a deformation that remapped the entire 3D mesh into a new global cartesian system. In this new system, the Z-axis represents the distance along the medial axis and the Y-axis is aligned with the medial axis to the dorsal vector.

This transformation effectively maps the neural tube into a cylindrical geometry while preserving all vertex-wise scalar data, including local curvature, tissue thickness, and proliferation density. Finally, the remapped 3D mesh was unwrapped into a 2D plane by projecting the coordinates into a cylindrical coordinate system (r, θ, z). The angular component (θ) and the longitudinal height (z) were extracted, with height subsequently normalized to a range between 0 and 1. This resulted in a standardized 2D map where local biological features could be visualized as heatmaps, enabling direct spatial comparisons across samples independent of their original 3D posture while accounting for variability in embryo size and developmental stage.

### Finite Element Modeling

To evaluate the mechanical response of the neuroepithelium under internal pressure, a 2D plane-stress finite element pipeline was developed using the Open-Source Python package SolidSpy (45). The cross-sectional geometry was parametrically defined by radial and thickness profiles derived from experimental measurements at the dorsal, lateral, and ventral anchor positions (Table S1). These parameters were continuous along the dorsoventral axis (0° to 180°) using a Rational Quadratic Bézier interpolation scheme with zero-slope boundary conditions at the poles to ensure *C*^1^ continuity. The domain was discretized into a structured mesh of four-node isoparametric quadrilateral elements (𝑛θ = 80, 𝑛𝑟 = 20). The tissue was modeled as an inert, linear elastic, and isotropic material with a Poisson’s ratio (ν) of 0.45 and a shear modulus (µ) of 300 Pa. A uniform normal internal pressure (P = 15 Pa) was applied as a follower load distributed as nodal forces along the inner boundary surface normal, while the outer boundary remained traction-free. Symmetry boundary conditions were imposed along the vertical axis (𝑢_𝑥_ = 0) at the 0° and 180° poles, with a single basal node pinned vertically (𝑢_𝑦_ = 0) to eliminate rigid body motion. Post-processing involved transforming the Cartesian stress tensor into a local polar basis to resolve transverse hoop stress (σ_θ_). To characterize the structural mechanics independent of absolute stiffness, all computed values were normalized by the shear modulus. The normalized average membrane stress (σ_𝑚𝑒𝑚_ /µ), representing the net tensile force per unit length, was continuously integrated across the wall thickness (t) according to: 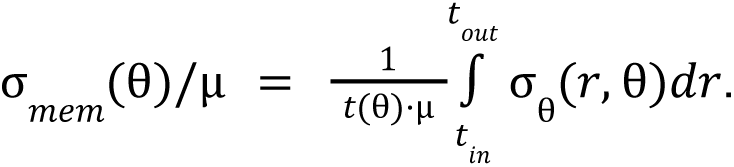 This enabled a direct correlation between spatial thickness gradients, localized bending-to-tensile structural inversions, and the cross-sectional geometry.

## Statistical Analysis

Group comparisons between developmental stages and compartments were performed using two-sample Welch’s t-tests (SciPy). Significance thresholds were defined as *p < 0.05, **p < 0.01, and ***p < 0.001. For correlations between tissue properties and curvature, data were pooled per stage and divided into ten equal-population quantile bins, and Pearson correlation coefficients were computed using scipy.stats.pearsonr. For cross-stage comparisons of dorso-ventral distributions, data were binned into 40 equal-width azimuthal bins spanning −180° to 180°, and Spearman correlation coefficients were computed using scipy.stats.spearmanr to assess conservation of spatial patterns between stages. All data are presented as mean ± standard deviation unless otherwise stated.

## Data and code availability

The custom Python scripts and computational meshes used to generate the results in this study, including 3D mesh processing, thin-plate spline warping, and 2D cylindrical mapping, are openly available on GitHub at https://github.com/nchahare/spatchcocking. These scripts utilize open-source libraries for geometric transformations and scalar field calculations. The raw datasets supporting the findings of this study are available from the corresponding author upon request.

## Author contributions

N.C. and N.N. conceived and designed the project. N.C. performed all experiments, including embryo preparation, immunostaining, cryosectioning, and confocal imaging. N.C. carried out image segmentation, 3D mesh reconstruction, morphometric analysis, and computational pipeline development. C.I. assisted with embryo preparation, cryosectioning, and image segmentation. N.C. and N.N. wrote the manuscript.

## Acknowledgments

We thank the members of the Nerurkar Lab as well as Giacomo Gattoni, PhD, for their scientific input and valuable feedback throughout this study. We further thank Marco Musy, PhD, for guidance regarding the vedo Python package. This work was funded by the National Institutes of Health (R35 GM142995 to N.L.N.).

---

**Figure S1.**
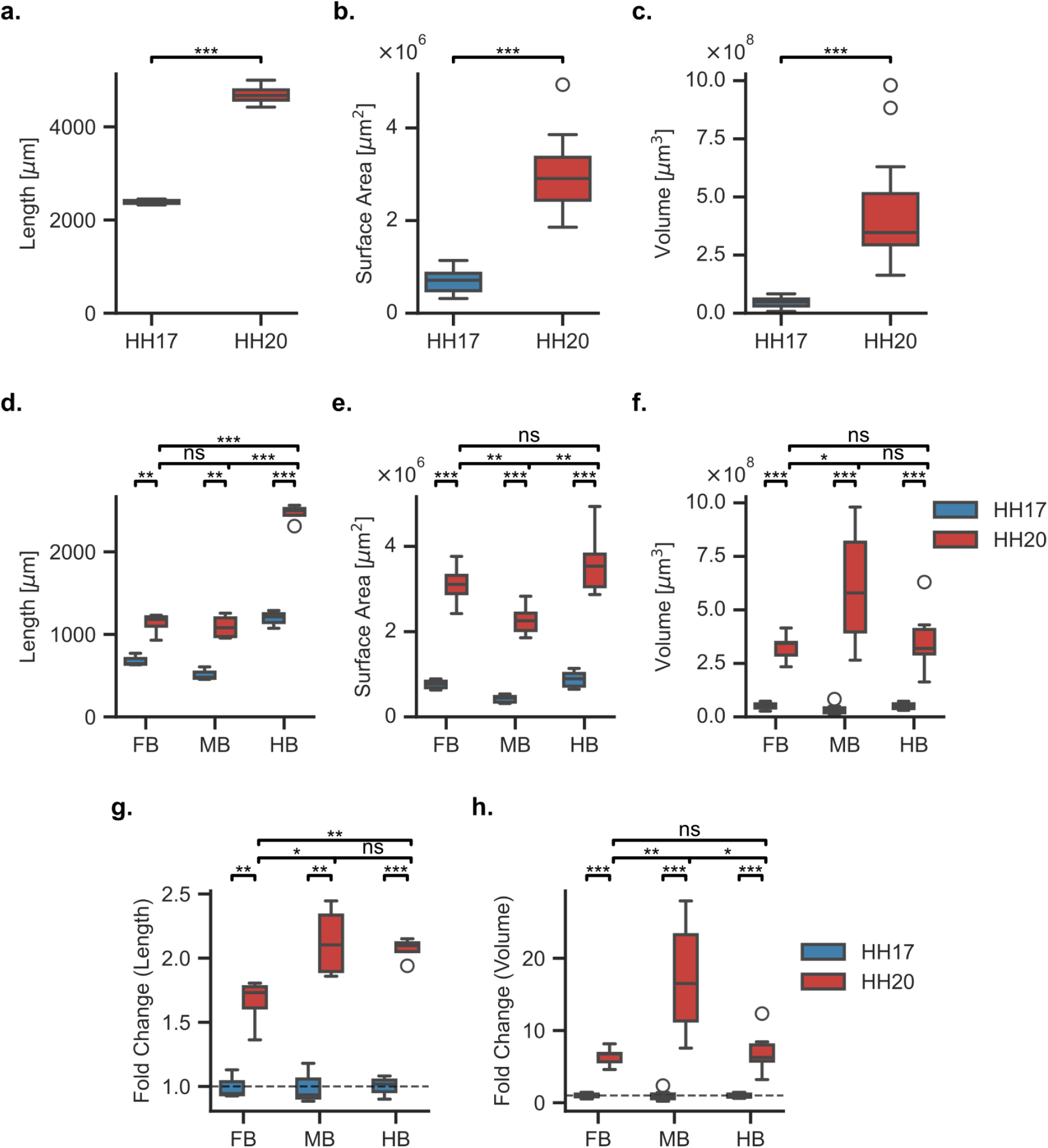
Regional morphometric measurements of cranial neural tube expansion between HH17 and HH20. (a–c) Lumen length (a), surface area (b), and volume (c) at HH17 and HH20. (d–f) Regional length (d), surface area (e), and volume (f) by primary vesicles at HH17 (blue) and HH20 (red). (g, h) Regional fold changes in length (g) and volume (h) by primary vesicles. Dashed line indicates fold change of 1. n = 7 embryos per stage. *p < 0.05, **p < 0.01, *p < 0.001, ns = not significant. FB, forebrain; MB, midbrain; HB, hindbrain.

**Figure S2.**
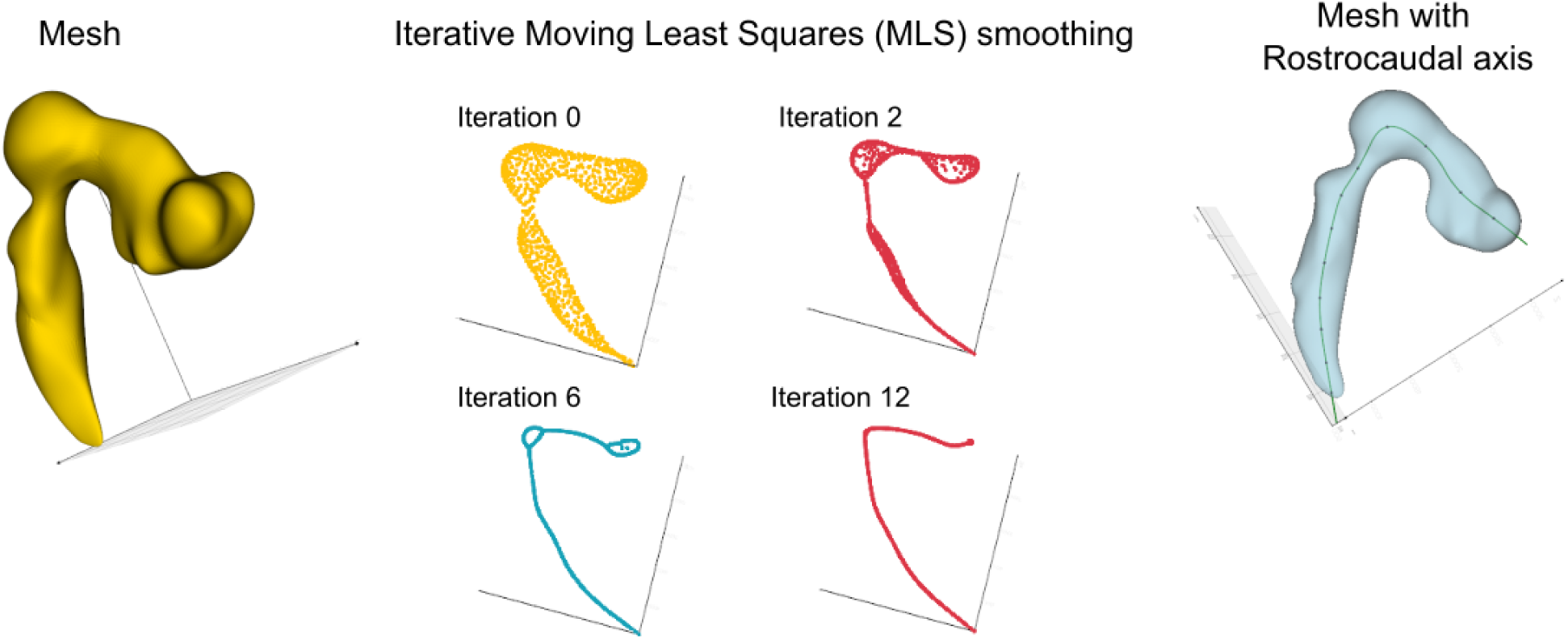
Medial axis extraction via iterative Moving Least Squares (MLS) smoothing. **(Left)** Triangular mesh of the cranial neural tube lumen used as input for medial axis extraction. **(Center)** Progressive iterations of MLS smoothing (iterations 0, 2, 6, and 12), showing convergence from a noisy point cloud to a smooth, stable medial axis curve tracing the rostral-caudal extent of the neural tube. **(Right)** Final extracted medial axis (rostrocaudal axis) overlaid on the lumen mesh, providing the reference frame for cross-sectional plane generation and the spatchcocking coordinate transformation.

**Figure S3.**
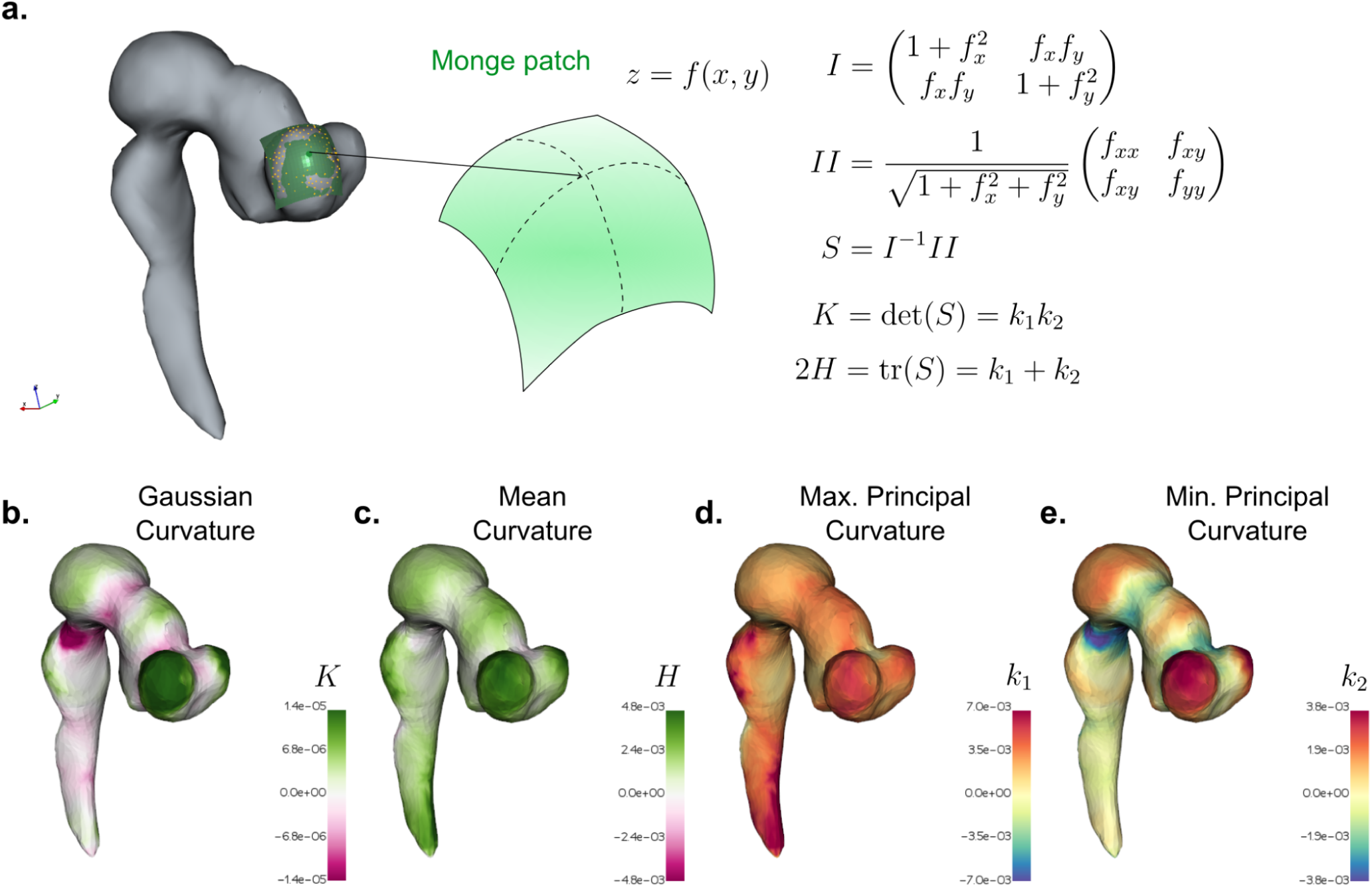
Vertex-wise curvature calculation via Monge patch fitting. **(a)** At each vertex of the basal mesh, a quadratic surface (Monge patch, z = f(x, y)) is fit to the local neighborhood. The first (I) and second (II) fundamental forms are derived from the partial derivatives of this surface, and the shape operator *S* = *I*⁻¹*II* yields Gaussian curvature (*K* = det(*S*) = *k*₁*k*₂) and mean curvature (2𝐻 = tr(*S*) = *k*₁ + *k*₂), where k₁ and k₂ are the principal curvatures. **(b–e)** Gaussian curvature 𝐾 (b), mean curvature 𝐻 (c), maximum principal curvature k₁ (d), and minimum principal curvature k₂ (e), mapped onto the basal surface of a representative HH20 embryo.

**Figure S4.**
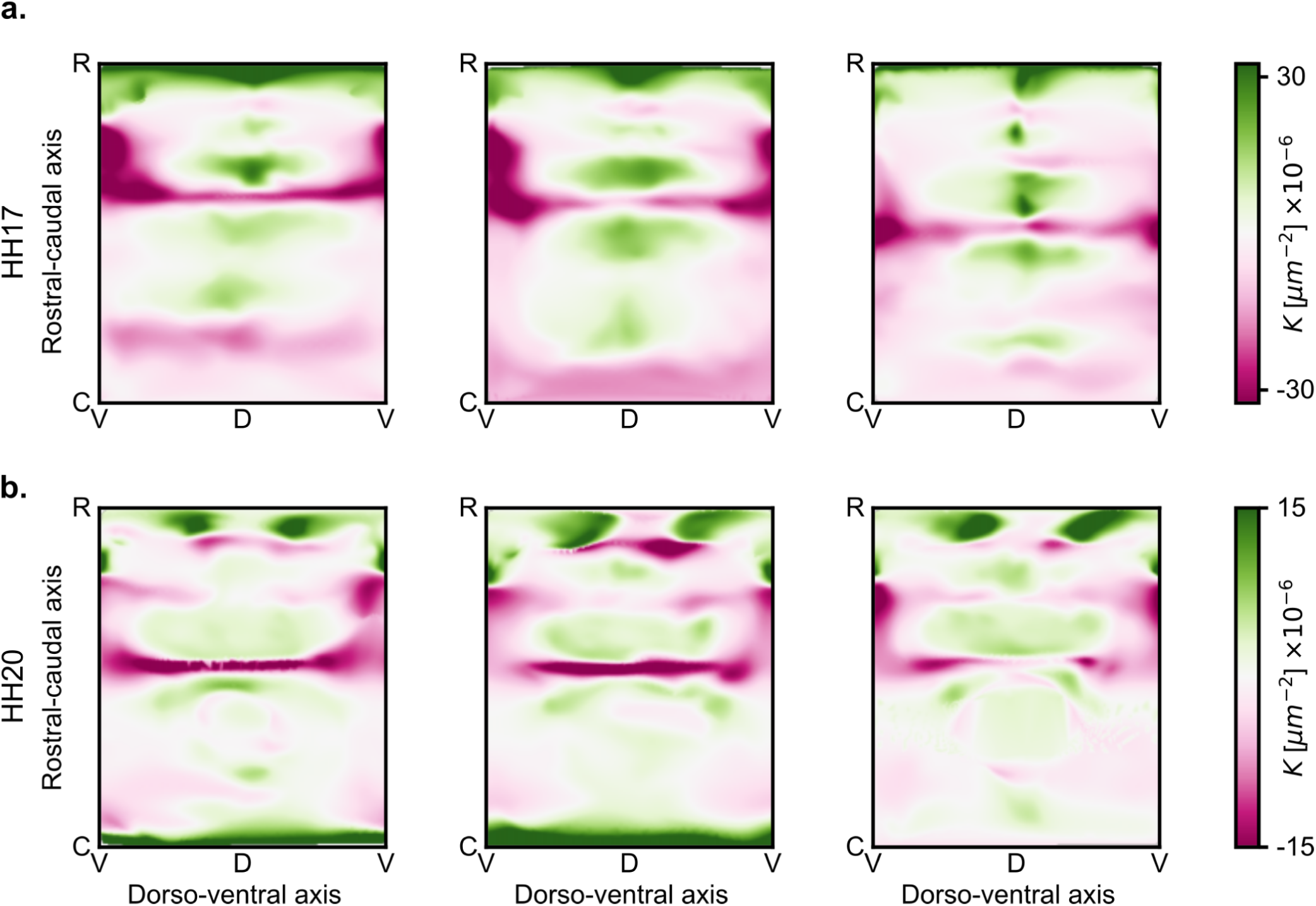
Gaussian curvature spatchcocked projections across individual embryos. **(a, b)** Spatchcocked 2D projections of Gaussian curvature (𝐾) for representative individual embryos at HH17 (top row) and HH20 (bottom row), mapped onto the standardized rostro-caudal and dorso-ventral coordinate system. Dorsal (D) midline is centered; ventral (V) is at either side. Color scales differ between stages. 𝐾, Gaussian curvature. Green indicates positive 𝐾; magenta indicates negative 𝐾.

**Figure S5.**
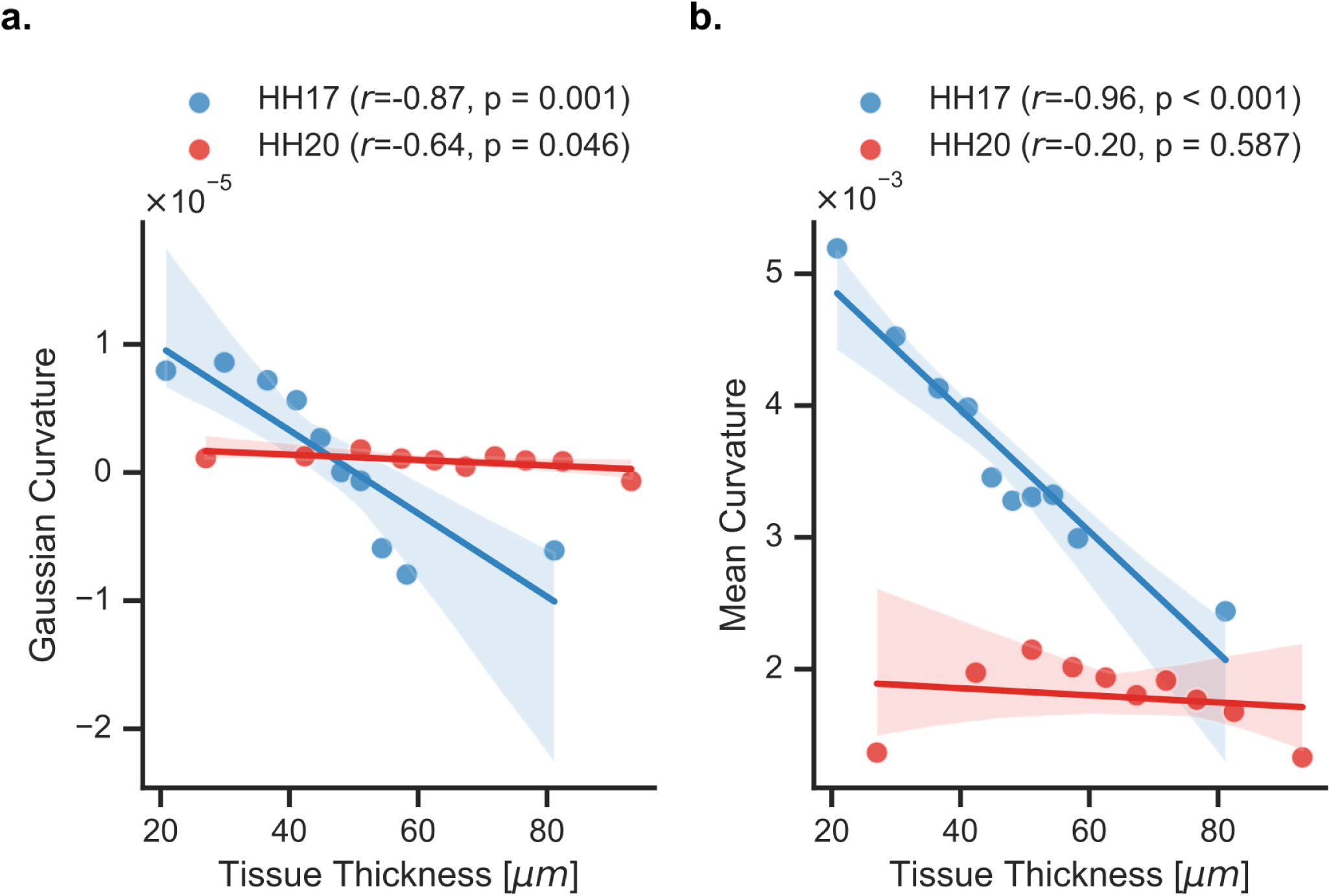
Decoupling of tissue thickness and surface curvature during expansion. **(a, b)** Scatter plots showing the relationship between tissue thickness and Gaussian curvature (a) and mean curvature (b) at HH17 (blue) and HH20 (red). Each point represents the mean of a quantile-defined bin (n = 10 equal-population bins) computed from pooled data across all embryos within each stage. Pearson correlation coefficients and associated p-values are reported for each stage. Shaded regions indicate 95% confidence intervals. K, Gaussian curvature; 𝐻, mean curvature. n = 4 embryos per stage.

**Figure S6.**
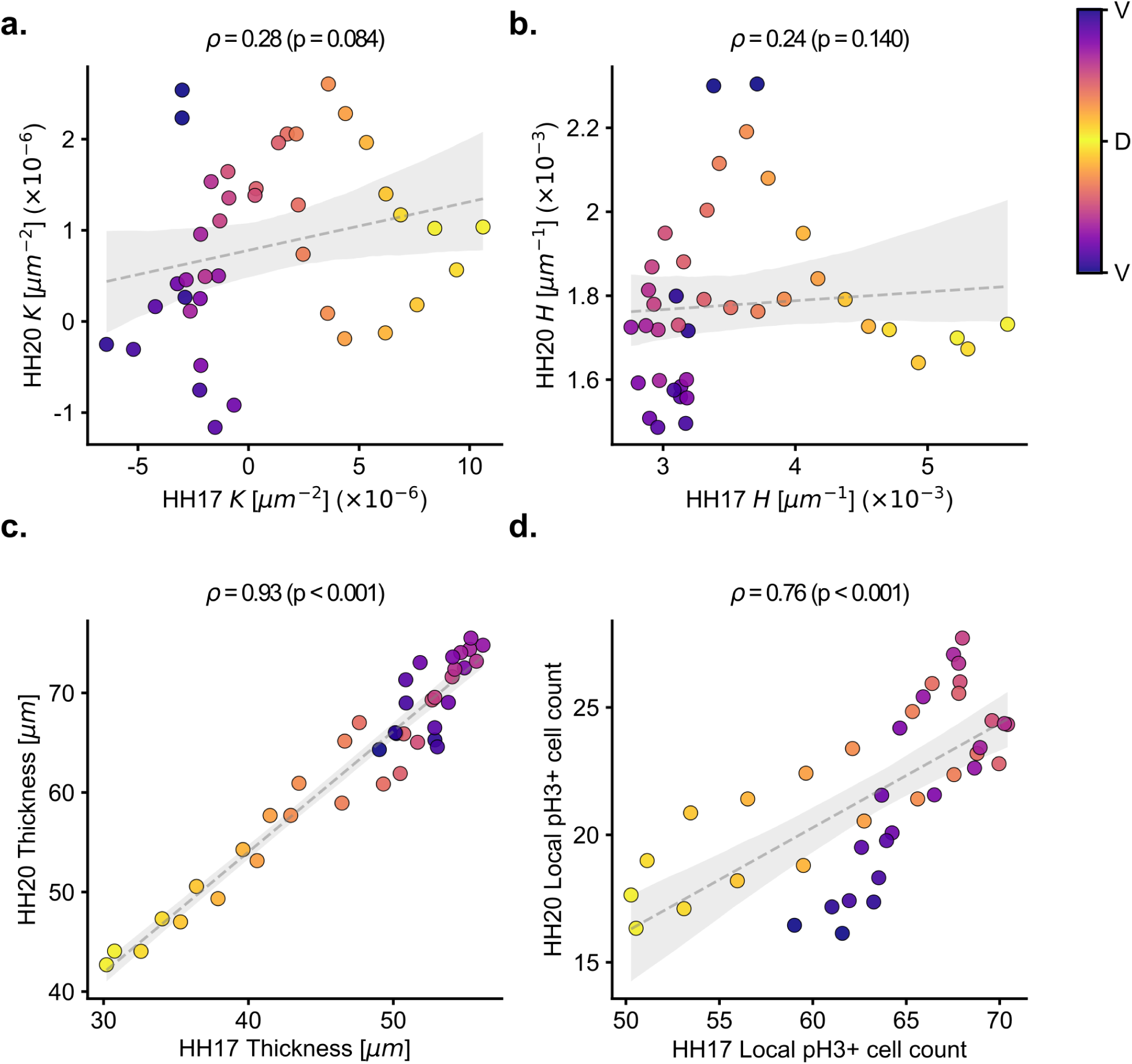
Dorso-ventral distributions of curvature, thickness, and mitotic activity are differentially conserved between HH17 and HH20. **(a–d)** Scatter plots comparing dorso-ventral distributions of Gaussian curvature 𝐾 (a), mean curvature 𝐻 (b), tissue thickness (c), and local pHH3+ cell count (d) between HH17 and HH20. Each point represents the mean value within a dorso-ventral angle bin (n = 40 equal-width bins spanning −180° to 180°), averaged across all embryos within each stage. Points are colored by dorso-ventral position. Spearman correlation coefficients and associated p-values are reported for each comparison. Shaded regions indicate 95% confidence intervals. 𝐾, Gaussian curvature; 𝐻, mean curvature; D, dorsal; V, ventral. n = 4 embryos per stage.

**Figure S7.**
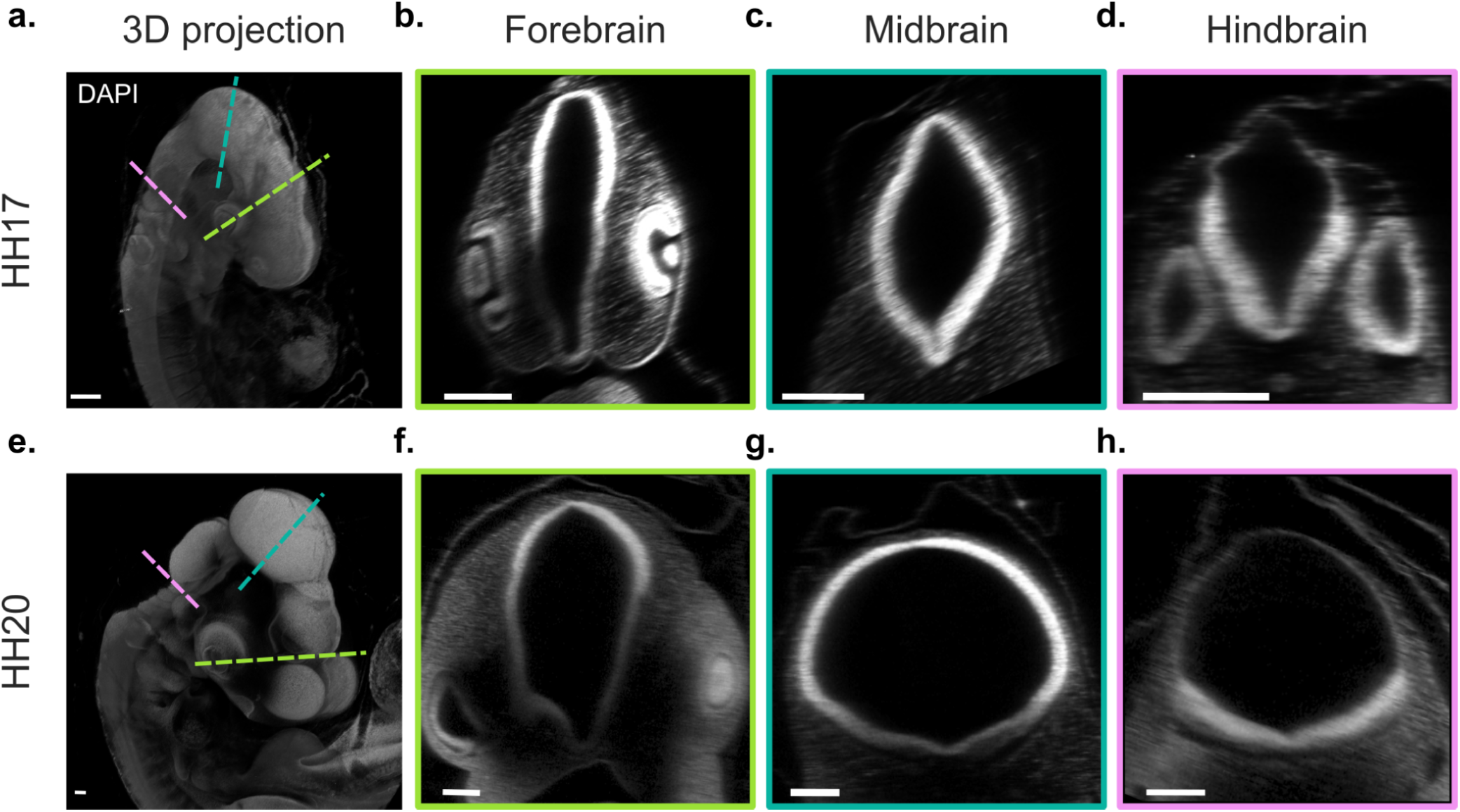
Cross-sectional anatomy of the cranial neural tube at HH17 and HH20. **(a, e)** Representative maximum intensity projections of cleared whole-mount embryos stained for DAPI at HH17 (a) and HH20 (e). Dashed lines indicate the planes of cross-sections shown in (b–d) and (f–h) for the forebrain (green), midbrain (cyan), and hindbrain (magenta). **(b–d, f–h)** Cross-sections through the forebrain (b, f), midbrain (c, g), and hindbrain (d, h) at HH17 (top row) and HH20 (bottom row). Scale bars, 200 µm.

**Figure S8.**
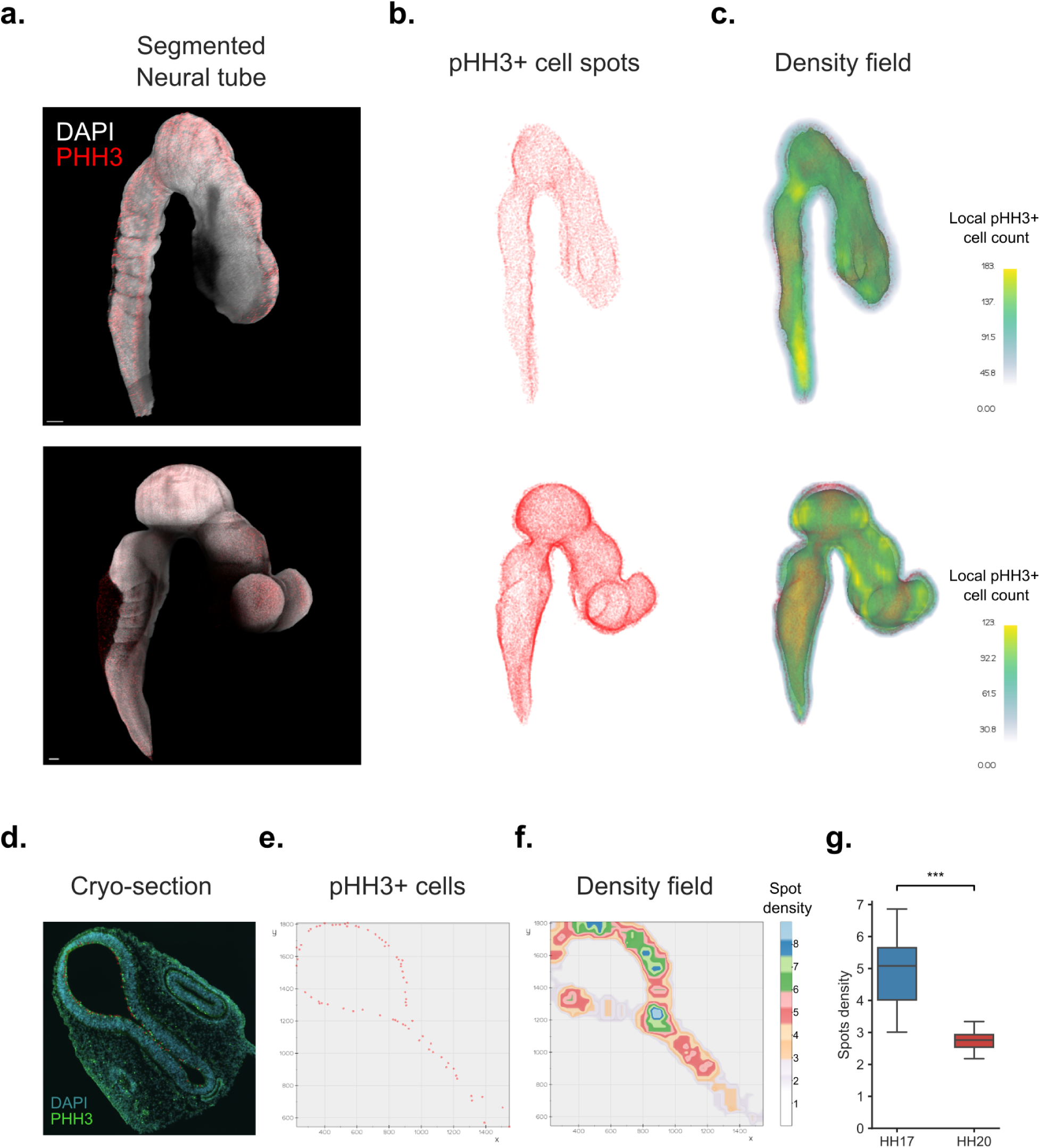
pHH3+ cell detection and density field computation pipeline. **(a)** Maximum intensity projections of representative HH17 (top) and HH20 (bottom) embryos stained for DAPI (white) and pH3 (red), with the segmented neural tube surface overlaid. Scale bars, 200 µm. **(b)** 3D point clouds of detected pHH3+ nuclei within the segmented neural tube at HH17 (top) and HH20 (bottom). **(c)** Local pHH3+ cell count mapped onto the 3D surface of the neural tube at HH17 (top) and HH20 (bottom), computed via fixed-radius neighborhood search (R = 100 µm) and transferred onto the basal mesh using inverse distance weighting interpolation. Note the difference in color scale between stages. **(d)** Representative cryosection stained for DAPI (cyan) and pH3 (green), illustrating the apical enrichment of pHH3+ nuclei within the neuroepithelium. **(e)** 2D point cloud of pHH3+ nuclei detected within the cryosection shown in (d). **(f)** Kernel density estimate of pHH3+ cell distribution from the cryosection shown in (e). **(g)** Mean Spot density of pHH3+ cells in cryosections at HH17 and HH20. ***p < 0.001. n = 4 embryos per stage.

**Figure S9.**
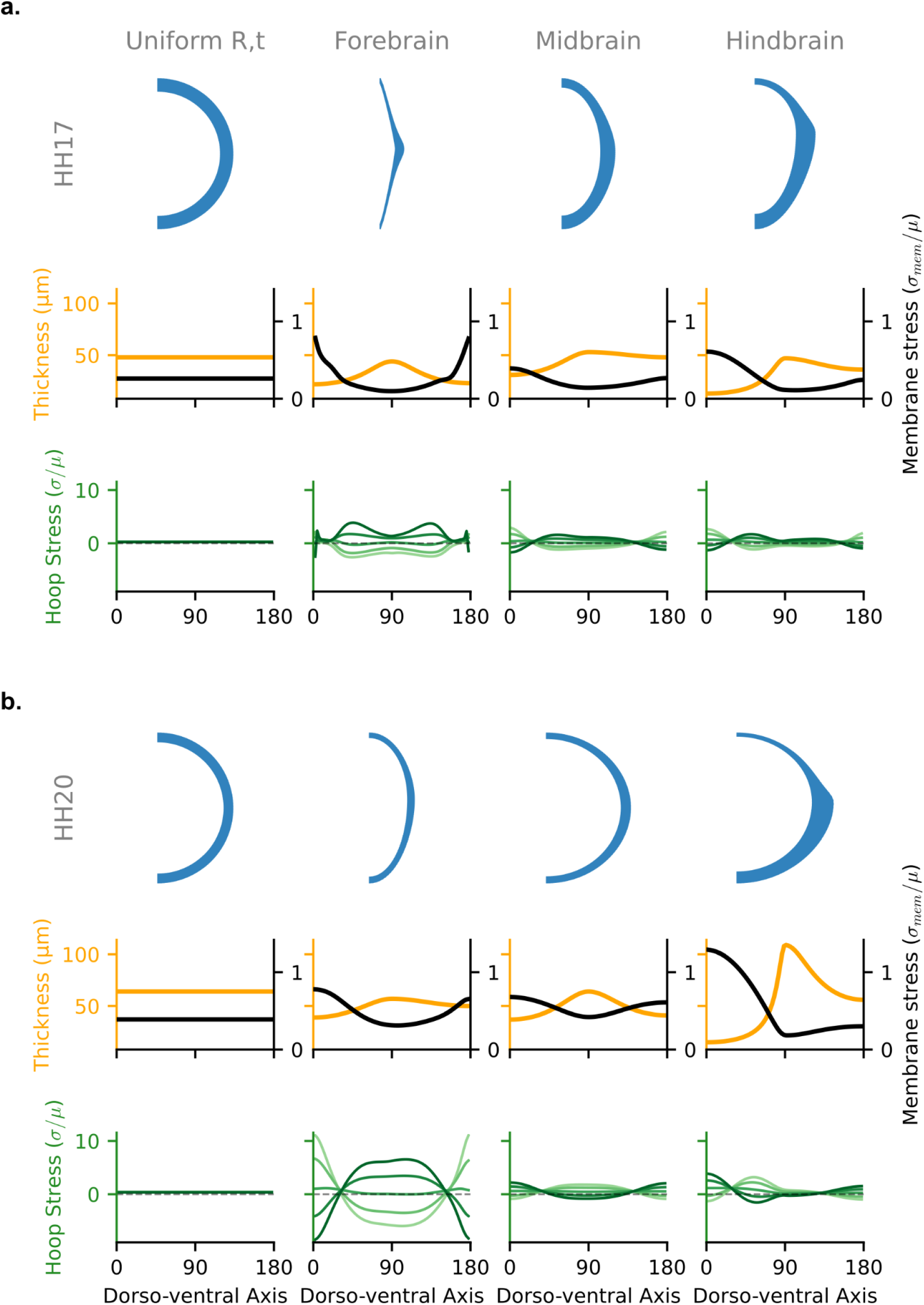
Plane-stress finite element modeling of the neuroepithelium under internal pressure. **(a)** HH17 and **(b)** HH20 cross-sectional simulations for 2-D cross sections of a shell of uniform thickness, and for geometries based on experimentally measured geometries of the forebrain, midbrain, and hindbrain. Each panel shows the modeled cross-sectional geometry (blue, top), thickness profile (yellow) and mean membrane stress (black) as a function of dorsoventral axis (middle), and hoop stress distributions at distinct apico-basal positions from inner/basal (dark green) to outer/apical (light green) (bottom). The finite element model assumes plane-stress conditions for a passive linear elastic solid with uniform internal pressure (P = 15 Pa) applied normal to the inner boundary; mediolateral symmetry is assumed and only the right hemicircle is modeled. All stress values are normalized by the shear modulus (σ/μ); membrane stress represents the thickness-integrated hoop stress per unit length (σ_mem_/μ), enabling comparison of structural mechanics independent of absolute material stiffness.

**Table S1:** Geometric input parameters and Bézier interpolation weights for finite element simulations. Input parameters, regional anchors, and interpolation weights per condition. The dataset tracks absolute inner dimensions (r), wall thickness metrics (t), and the corresponding Bezier interpolation shaping weights used by the FEA preprocessing script across the dorsal (top), lateral (mid), and ventral (bot) coordinates.

| Stage | Case | $r_{top}$ (μm) | $r_{mid}$ (μm) | $r_{bot}$ (μm) | $t_{top}$ (μm) | $t_{mid}$ (μm) | $t_{bot}$ (μm) | Radial Weight | Thickness Weight |
| --- | --- | --- | --- | --- | --- | --- | --- | --- | --- |
| HH17 | Uniform | 250 | 250 | 250 | 48 | 48 | 48 | 0.7071 | 0.7071 |
|  | Forebrain (FB) | 366 | 86 | 420 | 22 | 44 | 23 | 0.1 | 0.7071 |
|  | Midbrain (MB) | 250 | 150 | 250 | 31 | 53 | 48 | 0.7071 | 0.7071 |
|  | Hindbrain (HB) | 127 | 106 | 208 | 13 | 47 | 36 | 0.7071 | 0.7071 |
| HH20 | Uniform | 500 | 500 | 500 | 64 | 64 | 64 | 0.7071 | 0.7071 |
|  | Forebrain (FB) | 569 | 352 | 715 | 39 | 57 | 50 | 0.7071 | 0.7071 |
|  | Midbrain (MB) | 500 | 540 | 500 | 37 | 64 | 41 | 0.7071 | 0.7071 |
|  | Hindbrain (HB) | 351 | 400 | 371 | 15 | 109 | 56 | 0.7071 | 0.7071 |

